# Tau, amyloid-β and α-synuclein co-pathologies synergistically enhance neuroinflammation and neuropathology

**DOI:** 10.1101/2024.10.13.618101

**Authors:** Jhodi M. Webster, Ya-Ting Yang, Aidan T. Miller, Asta Zane, Kasandra Scholz, William J. Stone, Nikhita Mudium, Nicole J. Corbin-Stein, Woong-Jai Won, Anna C. Stoll, Kelsey M. Greathouse, Noelle H. Cooper, Lillian F. Long, Phaedra N. Manuel, Jeremy H. Herskowitz, Talene A. Yacoubian, Daniel J. Tyrrell, Ivette M. Sandoval, Fredric P. Manfredsson, Jeffrey H. Kordower, Ashley S. Harms

## Abstract

Alzheimer’s (AD) and Parkinson disease (PD) pathology often co-occur. Amyloid-β and phosphorylated tau are found in 30-50% of idiopathic PD cases, while α-synuclein inclusions are present in 50% of AD cases. These co-pathologies are linked to increased mortality and earlier onset of cognitive decline. Immune activation is a hallmark of these neurodegenerative diseases, but current models primarily examine each pathology in isolation. How these co-pathologies drive inflammation and neuronal loss remains poorly understood. We therefore developed a mouse model combining tau, amyloid-β, and α-synuclein. We found that co-pathologies synergistically trigger an amplified neuroimmune response, with expanded populations of CD4^+^ and CD8^+^ tissue-resident memory T cells and CD68^+^ microglia, compared to single pathologies. These changes were abundant in the hippocampus and cortex, regions with elevated protein pathology load and enhanced neuronal loss. Our findings demonstrate that co-pathologies enhance proteinopathy and synergistically enhance immune activation and neurodegeneration, suggesting that combinatorial therapeutic strategies that target both co-pathologies and inflammation, may be disease modifying.

**Summary:** Webster et al. demonstrate that co-occurring Alzheimer’s and Parkinson disease protein pathologies, common in cognitively impaired patient populations, amplify proteinopathy and synergistically enhance CNS neuroinflammatory responses and neurodegeneration. This work supports the need for combinatorial therapeutic strategies and positions neuroinflammation as an important link for co-pathology enhanced neurodegeneration.

## Introduction

In addition to the progressive loss of neurons and brain atrophy, neurodegenerative diseases share the common feature of pathogenic protein accumulation and worsening cognitive decline. These proteins include α-synuclein (α-syn) inclusions found in Lewy bodies and Lewy neurites in Parkinson disease (PD) and Parkinson disease dementia (PDD), extracellular amyloid-β (Αβ) plaques and phosphorylated-tau (tau) inclusions in the form of intracellular neurofibrillary tangles in Alzheimer’s disease (AD) (Goedert, 1993, Joachim and Selkoe, 1989, Spillantini et al., 1997). Although these proteins are pathologically associated with their respective diseases, there is evidence for the overlap of pathology and symptoms. The presence of these AD-related pathologies, found in 50-80% of PD brains (Irwin et al., 2017, Irwin et al., 2018, Jellinger, 2002, Jellinger, 2003, Jellinger, 2020, Kotzbauer et al., 2012, Robinson et al., 2018, Spires-Jones et al., 2017), has been a significant contributor to exacerbated disease progression and dementia onset in patients with PD (Compta et al., 2014, Kotzbauer et al., 2012). Similarly, Lewy bodies have been found in the brains of AD patients (Twohig and Nielsen, 2019, van der Gaag et al., 2024) and this a-syn co-pathology has been shown to promote amyloid-driven tau accumulation in AD brains (Franzmeier et al., 2025). The co-expression of these pathologies has been linked to earlier onset of cognitive decline and higher rate of mortality (Coughlin et al., 2019, Kraybill et al., 2005, Mak et al., 2024, Malek-Ahmadi et al., 2019). This exacerbated disease phenotype under co-pathology conditions suggests potential synergy between α-syn, tau and Aβ, however the disease-progressing mechanisms of this co-expression remain unknown.

Previously, multi-pathology mouse models have also implicated potential interactions between pathologies. In dementia with Lewy body (DLB)-AD and PD-AD mouse models, combined tau, Aβ and α-syn pathologies enhanced aggregation and spreading of pathology beyond single pathology effects (Bassil et al., 2020, Bassil et al., 2021, Clinton et al., 2010). Neurodegeneration, however, was only shown to be influenced by Aβ and α-syn co-pathology in 5XFAD mice injected with α-syn mouse pre-formed fibrils (PFFs) (Bassil et al., 2020). Additionally, neuroinflammation in the presence of the co-pathologies was not fully assessed in these animal models. As such, it remains unknown how these co-pathologies may promote the immune cell activation leading to the neurodegeneration and sustained aggregated protein pathology seen in human disease.

Neuroinflammation, driven by microglia and CNS-resident macrophages, infiltrating lymphocytes, and monocytes, is a key contributor to neuronal loss and neurodegenerative disease progression. CD4^+^ and CD8^+^ T cells are found in the brains (Itagaki et al., 1988, Merlini et al., 2018, Rogers et al., 1988, Togo et al., 2002) and cerebrospinal fluid (Gate et al., 2020) of AD, PD and DLB patients (Brochard et al., 2009, Sulzer et al., 2017, Yan et al., 2021), along with activation of myeloid cells, including microglia, macrophages and monocytes. In AD, Aβ plaques trigger and can interact with microglia to adopt a phagocytic but dysfunctional disease-associated microglia (DAM) phenotype (Ries and Sastre, 2016, Zhao et al., 2018) while Aβ-specific T cells infiltrate the brain and exacerbate tau pathology and neurodegeneration (Keren-Shaul et al., 2017). Tau pathology also triggers a pro-inflammatory phenotype in microglia through receptor binding (Pampuscenko et al., 2023) and promotes CNS-specific changes in T cell reactivity (Chen et al., 2023). In PD, α-syn also interacts with toll-like receptors (TLRs) on microglia to promote release of inflammatory cytokines (Karpenko et al., 2018), while T cells target α-syn epitopes in the substantia nigra pars compacta (SNpc) (Ma et al., 2025, Sulzer et al., 2017). While these cells aid in immune surveillance during homeostatic conditions, in chronic disease, microglia and T cells may adopt pro-inflammatory phenotypes and contribute to neurotoxicity and exacerbated neuronal damage (Altendorfer et al., 2022, Galiano-Landeira et al., 2020, Hefendehl et al., 2014, Kimura et al., 2024, Wang et al., 2021). Although recent work has highlighted enhanced microglial activation in mixed DLB + AD patients (van Wetering et al., 2024), how combined tau, Aβ and α-syn pathologies may synergistically influence the full neuroinflammatory response remains unclear.

To further understand how co-pathologies drive neurodegeneration, we aimed to determine whether the combined presence of tau, Αβ and α-syn promotes neuroinflammation, protein aggregation and neuronal loss through synergistic effects. To do this, we developed a co-pathology mouse model, generated by injection of adeno-associated virus (AAV)-double mutant tau into the entorhinal cortex and α-syn mouse PFFs into the striatum of J20 amyloid over-expressing transgenic mice (Mucke et al., 2000). This model allowed us to evaluate the impact of multiple pathological proteins on innate and adaptive immune responses as well as neurodegeneration within vulnerable regions, including the hippocampus. We found that the presence of co-pathologies not only promotes protein pathology deposition compared to single pathology models, but also triggered synergistic neuroinflammatory responses and subsequent enhanced neuronal loss in the hippocampus. These findings support post-mortem co-pathology data (van Wetering et al., 2024) and model tau, Aβ and α-syn synergizing to enhance key immunopathogenic processes that may contribute to neurodegenerative disease progression.

## Results

### Co-pathologies enhance Aβ plaques and α-syn inclusions

The co-expression of tau, Aβ and α-syn has been reported in the limbic and cortical regions, particularly in the hippocampus and neocortex in human postmortem studies from DLB, AD and PDD brains (Colom-Cadena et al., 2013, Ferman et al., 2018, Kouli et al., 2020). To model this *in vivo* in mice, we generated a co-pathology model incorporating tau, Aβ and α-syn and compared data obtained in this co-pathology model with mice only expressing a single pathology. As a Tau single pathology control, tau pathology was induced in non-transgenic (NTG), C57BL/6J mice via bilateral entorhinal cortex injection of an AAV expressing human tau with two mutations (AAV9-Tau) (Figure 1a). To model Aβ single pathology, J20 transgenic (Tg) mice (Mucke et al., 2000), which overexpress human mutant amyloid precursor protein (APP^mut^) and develop Aβ plaques in the hippocampus and cortex (Figure 1b), were aged to 6 months old. Lastly, for α-syn single pathology control, 12–14-week-old NTG mice received bilateral injections of mouse α-syn PFFs into the dorsal lateral striatum (Figure 1c). To create a model of co-pathology, age-matched J20 mice were bilaterally injected with both α-syn PFFs into the basal lateral striatum and AAV9-Tau into the entorhinal cortex during one stereotaxic surgery session, combining tau, Aβ and α-syn pathologies in one model (Figure 1d) resulting in co-pathologies in the hippocampus and cortex.

**Figure 1.**
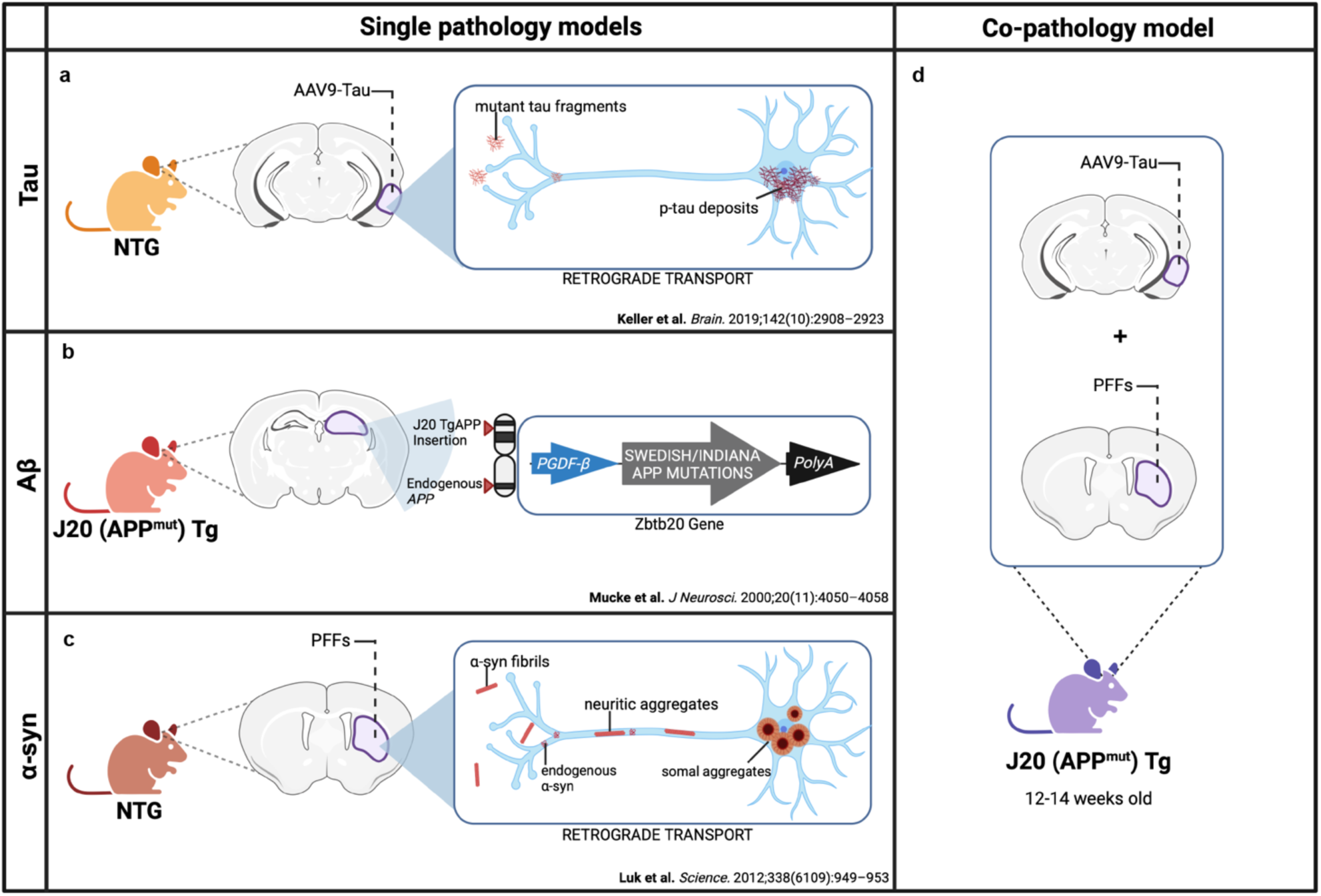
Strategy for creating co-pathology mouse model. At 12-14 weeks old, non-transgenic (NTG) wild-type or J20 APP mutant (APP^mut^) transgenic (Tg) age-matched males and females are used to induce either single pathology controls or the co-pathology model *in vivo*. (a) For tau single pathology controls, NTG mice received injections of 2μL of 3.2E12 vg/mL AAV9-Tau into the entorhinal cortex. This viral overexpression of mutant tau results in the accumulation of phosphorylated tau (p-tau) in the cell body via retrograde transport. (b) For Aβ pathology, J20 Tg mice expressing human APP with Swedish and Indiana mutations under the PGDF-β promoter were used. These mice develop Aβ plaques due to overproduction and deposition of Aβ. (c) For α-syn pathology in the PFF single pathology control, NTG mice were injected with 2μL of 5μg α-syn PFFs into the striatum. Fibrils deposited into the brain will interact with endogenous α-syn within the neurons to form neuritic and somal aggregates. (d) For the co-pathology model, J20 Tg mice received both AAV9-Tau and α-syn PFFs into the entorhinal cortex and striatum, respectively, to simultaneously induce the expression of Aβ, tau and α-syn the brain.

Co-pathologies exhibit synergistic interactions that promote the aggregation and accumulation of disease-associated protein (Franzmeier et al., 2025, Irwin et al., 2013, Robinson et al., 2018, Sengupta and Kayed, 2022). To determine whether the presence of co-pathology enhances α-syn, tau and Aβ individually, single pathology controls and co-pathology brains were assessed for histological and biochemical markers of protein pathology at 3- and 6-months post-induction (MPI) using immunohistochemistry (IHC), western blotting, and enzyme linked immunosorbent assay (ELISA) (Figure 2a). AT-8^+^ neurons were immunolabeled in the cortex and hippocampus of 6-month-old (3MPI) (Figure 2b) and 9-month-old (6MPI) (Supplemental Figure 1a) Tau single pathology and co-pathology mice. Total tau protein expression at 3MPI and 6MPI in Triton X-100 soluble and insoluble fractions of the hippocampus and cortex was assessed using western blot (Figure 2c). No significant difference in tau pathology was observed between Tau single pathology and co-pathology groups at 3MPI, but a significant difference in tau accumulation was observed at 6MPI. At 6MPI, the presence of co-pathologies enhanced soluble tau in the hippocampus of co-pathology mice compared to single pathology controls (Figure 2d). Collectively, these data suggest that tau pathology accumulates in the hippocampus and cortex over time, and highlight the presence of co-pathology enhanced soluble tau in the hippocampus of mice at 6MPI.

**Figure 2.**
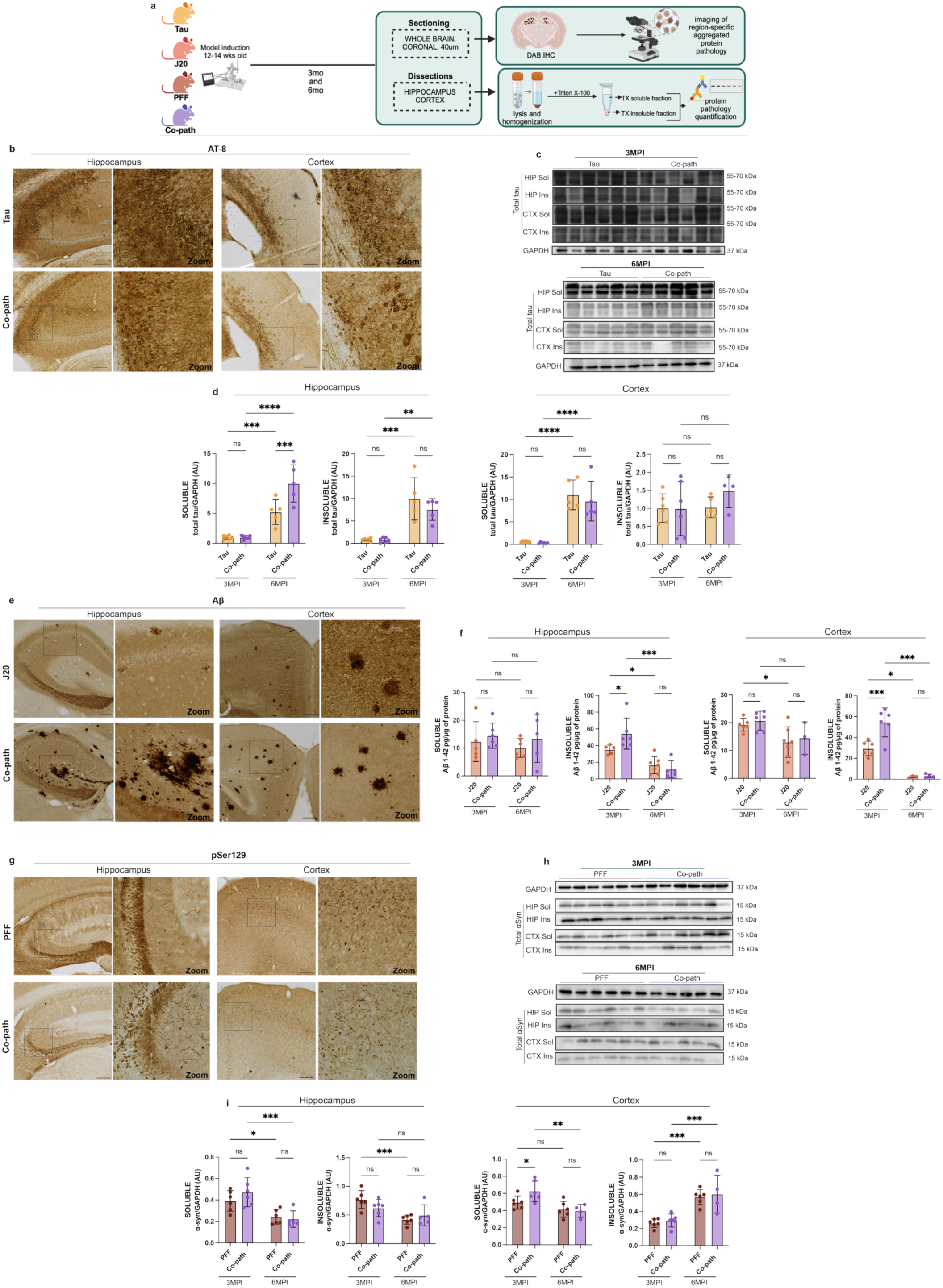
The presence of co-pathology enhances Aβ_1-42_ and α-syn protein levels in the hippocampus and cortex at 3MPI and total tau protein in the hippocampus at 6MPI. (a) 12–14-week-old animals were induced with either single pathologies or co-pathologies. At 3MPI and 6MPI, mice were sacrificed and brains processed for IHC or biochemical analysis of protein levels. (b) AT-8/pTau-positive neurons (DAB+, brown) in the hippocampus and cortex of tau single pathology and co-pathology brains. (c) Representative immunoblot of total tau and GAPDH levels from 3MPI and 6MPI co-pathology and Tau single pathology brains. Blots are cropped from original images found in Source Data. (d) Quantification of tau protein levels normalized to GAPDH levels in Triton X-100 (TX)-soluble and TX-insoluble hippocampus and cortical samples from Tau single pathology and co-pathology mice at 3MPI and 6MPI. (e) Aβ-positive plaques (DAB+, brown) expressed within the hippocampus and cortex of J20 single pathology (Aβ) and co-pathology brains. (f) ELISA was used to quantify Aβ_1-42_ levels in the brains of 3MPI and 6MPI J20 single pathology and co-pathology mice. TX soluble and insoluble fractions of 3MPI and 6MPI co-pathology and single pathology hippocampal and cortical tissue were used. (g) pSer129-positive inclusions in the hippocampus and cortex of PFF only (α-syn) and co-pathology brains. (h) Representative immunoblot of total α-syn and GAPDH. Blots are cropped from original images found in Source Data. (i) Quantification of synuclein protein levels normalized to GAPDH levels in TX-soluble and TX-insoluble hippocampus and cortical samples from PFF single pathology and co-pathology mice at 3MPI and 6MPI. All images taken at 10X or 20X for zooms, scale bars = 100μm. Analyzed using Welch’s t-test. *p<0.05, **p<0.01, ***p<0.001, ****p<0.0001. n=5-6 animals per group, both males and females used.

Using an antibody specific to Aβ aggregation, Aβ^+^ plaques were observed via immunohistochemistry in the hippocampus and cortex of co-pathology and J20 (Aβ) single pathology controls at 3MPI (Figure 2e) and 6MPI (Supplemental Figure 1b). As assessed by ELISA, insoluble Aβ_1-42_ protein decreased between 3MPI and 6MPI. At 3MPI, insoluble Aβ_1-42_ levels were significantly increased in the hippocampus and cortex of co-pathology brains compared to J20 single pathology controls indicating the presence of co-pathologies enhanced early insoluble Aβ_1-42_ (Figure 2f). pSer129^+^ α-syn inclusions were visible within the neurites and cell bodies in the hippocampus and cortex of PFF (α-syn) single pathology and co-pathology mice at 3MPI (Figure 2g) and 6MPI (Supplemental Figure 1c). Total α-syn protein expression at 3MPI and 6MPI in the hippocampus and cortex was assessed via western blot (Figure 2h). Similar to Aβ_1-42_, total α-syn protein was decreased in the hippocampus and cortex at 6MPI, compared to 3MPI. At 3MPI, there was an increase in soluble α-syn in the cortex of co-pathology brains compared to PFF single pathology (Figure 2i) controls.

All together, these results highlight that tau, Aβ and α-syn as co-pathologies may enhance accumulation of one another indicating that the location and timing of pathology deposition may be an important factor.

### Neuronal loss is driven by the co-expression of tau, Aβ and α-syn pathologies

Neurodegeneration, including changes in synaptic density and the subsequent death of neurons, occurs prior to severe onset of symptoms associated with neurodegenerative diseases. Such symptoms include cognitive and motor impairments (Braak et al., 2003, DeKosky and Scheff, 1990, Greffard et al., 2006, Selkoe, 2002). Given the involvement of tau, Aβ and α-syn in neurodegenerative diseases and their strongly association with these symptoms, we performed open field, Barnes maze, rotarod and pole behavioral tests to assess for cognitive and motor deficits in the model at 6MPI (Supplemental Figure 2). Sham mice, NTG animals bilaterally injected with α-syn monomer and AAV9-GFP into the striatum and entorhinal cortex, respectively, were used as a surgical control for co-pathology mice. Behavioral changes were observed with co-pathology mice making significantly more errors and reduced time to commit errors compared to sham mice in the Barnes maze test, but no significant difference was observed when compared to single pathology controls despite a faster time to the goal quadrant compared to J20 mice (Supplemental Figure 2a). Co-pathology mice showed increased turnaround and total time on the pole test compared to sham mice (Supplemental Figure 2b). The open field test also showed increased distance traveled by the co-pathology mice compared to sham (Supplemental Figure 2c). However, overall co-pathology mice showed no differences in movement as shown by the rotarod test (Supplemental Figure 2d). In summary, co-pathologies did not produce any major effects on motor or cognitive function in mice. This may be due to the intrinsic neural hyperexcitability of the J20 mouse model, which is known to drive a hyperactive behavioral phenotype (Busche et al., 2012, Shabir et al., 2020).

Dendritic spines are important for learning and memory and their morphology is an indicator of synaptic integrity and plasticity, both of which are disrupted in neurodegenerative diseases associated with cognitive decline (Bourne and Harris, 2008, Knobloch and Mansuy, 2008, Spires-Jones and Hyman, 2014). To test the hypothesis that tau, Aβ and α-syn co-pathologies synergize to exacerbate synaptic damage, we performed a detailed morphometric analysis of dendritic spines on Lucifer yellow labelled CA1 pyramidal neurons in the hippocampus across single and co-pathology brains. Our analysis showed that co-pathologies are associated with distinct changes to spine structure in the hippocampus. Specifically, co-pathology spines were significantly shorter in length compared to α-syn single pathology (Supplemental Figure 3a,b). Interestingly, spine volume on basal dendrites was increased in the hippocampus of co-pathology mice relative to brains with only Aβ or α-syn pathology, which may indicate a compensatory but dysfunctional state (Supplemental Figure 3c). Collectively, these findings suggest the presence of co-pathologies alters synapse integrity, which may be important for driving neuronal loss and cognitive decline.

Neuronal loss is a hallmark feature observed in both AD and PD, with region-specific vulnerability depending on disease pathology (Double, 2012, Giguère et al., 2018, Mrdjen et al., 2019, Olajide et al., 2021). Previous work, however, has only assessed neuronal loss in the context of α-syn and Aβ as co-pathologies (Bassil et al., 2021). Given this and our synaptic morphology findings, we next hypothesized that co-expression of tau, Aβ and α-syn may enhance neuronal loss in the hippocampus. To assess neurodegeneration, we performed unbiased stereology on neurons in the hippocampus of co-pathology and single pathology controls at 6MPI (Figure 3a). Immunostaining for NeuN+ neurons in the hippocampus of single pathology and co-pathology mice showed a distinct loss of neurons in the CA1 region of co-pathology brains (Figure 3b). Although there was an expected loss of neurons in the CA1 or CA3 regions of the hippocampus of each of the single pathology brains compared to their controls (Supplemental Figure 4a,b), we found enhanced loss (20-30%, p<0.001) of NeuN^+^ neurons in the CA1 region of the hippocampus in the co-pathology mice compared to tau, Aβ or α-syn single pathology mice (Figure 3c). Within the CA3, co-pathology mice showed significant loss (20-25%, p<0.01) compared to single tau and α-syn controls only (Figure 3d). Along with cognitive decline, patients with neurodegenerative disease also exhibit significant motor deficits, mostly due to the loss of neurons in motor-related cortical regions or loss of dopaminergic neurons in the SNpc (Braak et al., 2003, Buchman and Bennett, 2011). In the SNpc at 6MPI, the loss of tyrosine hydroxylase (TH)-positive neurons was significantly enhanced in the co-pathology model compared to that in the Aβ single pathology animals. However, in comparison to tau and α-syn mice, the co-pathologies did not enhance TH+ cell loss (Supplemental Figure 4c,d). Taken together, these results highlight a synergistic interaction between tau, Aβ and α-syn co-pathologies that exacerbates neurodegeneration in the hippocampus.

**Figure 3.**
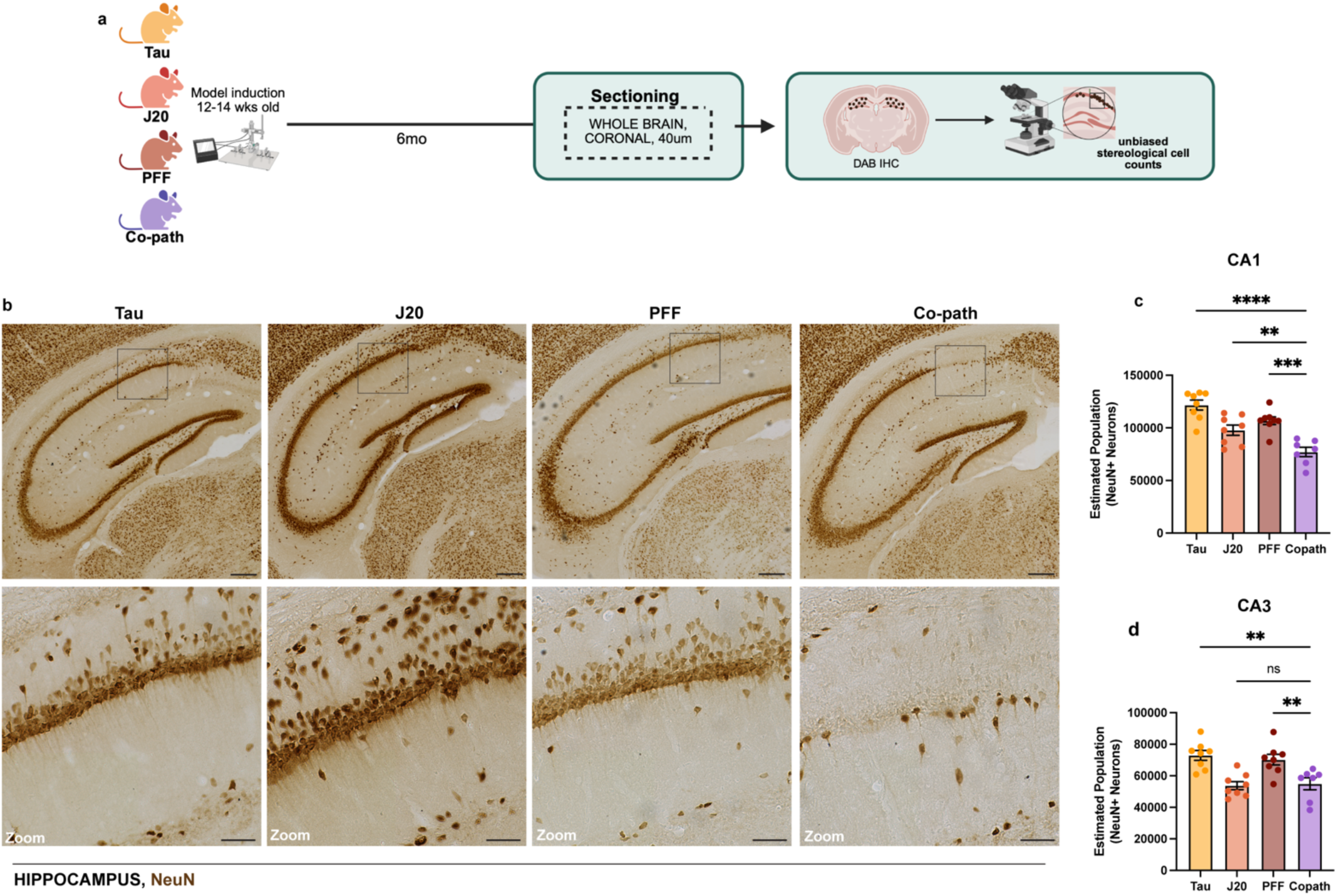
Co-pathologies drive enhanced neuronal loss in the hippocampus at 6MPI. (a) At 6MPI single pathology and co-pathology brains were processed for DAB IHC and unbiased stereology. (b) Representative brightfield images of NeuN+ neurons (DAB+, brown) in the hippocampus of Tau, J20 and PFF single pathology and co-pathology brain. Unbiased stereology was used to quantify the estimated number of NeuN+ neurons in the (c) CA1 and (d) CA3 regions of the hippocampus in the co-pathology mouse model compared to Tau, J20 and PFF single pathology brains. All images taken at 20X. Scale bars = 300μm and 100μm for zooms. One-way ANOVA with post hoc test was used to assess for significance. Mean values +/- SEM are plotted. ns = no significance, **p<0.01, ***p<0.005, ****p<0.0001. n=7-8 animals per group, both males and females used.

### Co-pathologies enhance microglial activation and infiltration of peripheral monocytes

Given that tau, Aβ and α-syn pathologies are individually associated with microglia activation and infiltration of peripheral myeloid cells in *in vivo* models (Harms et al., 2018, Heneka et al., 2013, Maphis et al., 2015) and in human disease(van Wetering et al., 2024), we aimed to determine the role of co-pathologies in enhancing innate immune responses in the CNS. Co-pathology and single pathology models were induced as previously stated using 12–14-week-old J20 Tg or NTG littermates (Figure 1). At 3MPI, hippocampal and cortical regions were dissected from co-pathology and single pathology controls for IHC analysis, mononuclear cell isolation and flow cytometry (Figure 4a). Using fluorescent IHC we immunolabeled Iba1, a marker used to visualize morphology of both homeostatic and activated microglia, in single pathology and co-pathology brains. Compared to tau, Aβ and α-syn single pathology brains, an increased number of “activated” Iba1^+^ cells were observed in the hippocampus of co-pathology mice (Figure 4b). Using flow cytometry, CD11b^+^CD45^+^ double positive myeloid cells were gated to analyze CNS resident microglia (CD45^mid^CX3CR1^+^) and CD45^hi^ CNS macrophage populations (including infiltrating monocytes and other CNS resident macrophages) (Supplemental Figure 5a; Figure 4c). In the hippocampus and cortex, the number of microglia and CD45^hi^ macrophages increased with co-pathologies (Figure 4d,e) compared to single pathology controls. Within this CD45^hi^ population, Ly6C^hi^ infiltrating monocytes and CNS resident macrophages were also increased in the hippocampus of co-pathology brains compared to tau, Aβ and α-syn single pathology brains (Supplemental Figure 5d,e).

**Figure 4.**
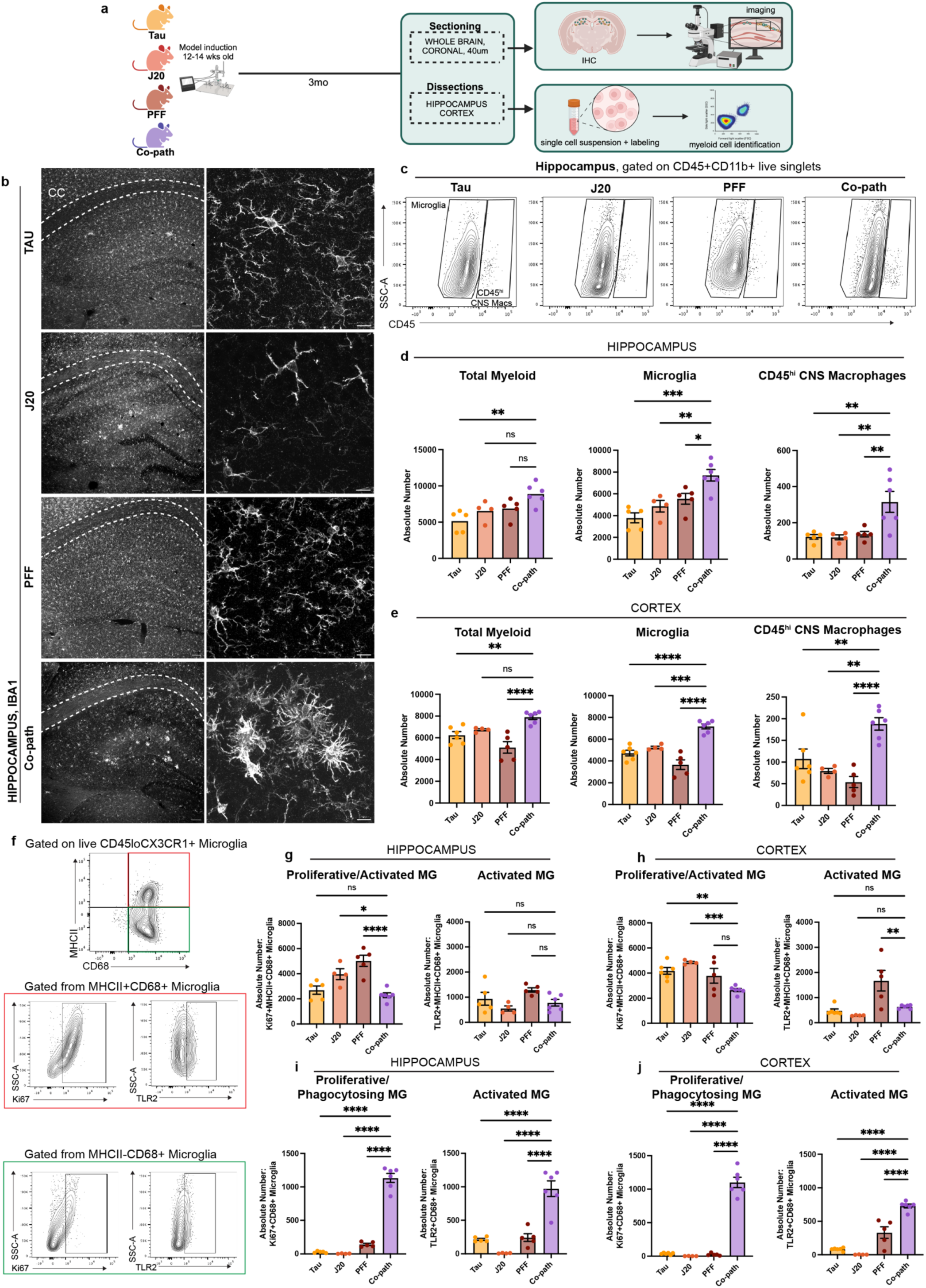
Co-pathologies enhance microglia activation at 3MPI. (a) J20 transgenic mice are injected at 12-14 weeks of age with 2uL of 5ug α-syn PFFs and 2uL of 3.2E12 vg/mL AAV9-Tau into the striatum and entorhinal cortex, respectively. For single pathology controls, age-matched non-transgenic littermates are injected with α-syn PFFs into the dorsal lateral striatum for PFF single pathology control or AAV9-Tau into the entorhinal cortex for Tau single pathology control. Age-matched J20 mice are used as the Aβ single pathology controls. At 3MPI, brains were collected and used for IHC or the hippocampus and surrounding cortices dissected and processed for spectral flow cytometry. (b) Representative fluorescent grayscale images of Iba1+ microglia in the hippocampus of Tau, J20 and PFF single pathology controls compared to that in the co-pathology model. (c) Representative flow cytometry contour plots showing CD45^mid^ microglia and CD45^hi^ CNS macrophages in single pathology and co-pathology hippocampus. The absolute number of total myeloid cells, microglia and CD45^hi^ infiltrating monocytes and resident macrophages in the (d) hippocampus and (e) cortex of Tau, J20 and PFF only compared to co-pathology brains are plotted. (f) From the CD45^mid^ microglia population, representative contour plots show expression of activation markers MHCII (antigen presentation, activated), CD68 (phagocytosing). The absolute number of MHCII^+^CD68^+^ microglia that are Ki67^+^ or TLR2^+^ in the (g) hippocampus and (h) cortex of co-pathology and single pathology mice are plotted. MHCII^-^CD68^+^ microglia that are Ki67^+^ or TLR2^+^ in the (i) hippocampus and (j) cortex of Tau, J20 and PFF only compared to co-pathology brains are plotted. Analyzed using One-way ANOVA. Mean values +/- SEM are plotted. *p<0.05, **p<0.01, ***p<0.005. n=5-6/group, with two brain regions pooled per sampled, both males and females included in analysis. First panel images taken at 10X magnification and zoomed, z-stack images at 40X. Scale bars (10X) = 100μm, scale bars (40X) = 10μm.

Given that microglia exhibit heterogeneity in health and disease based on their activation state, we next profiled markers of proliferation and activation to define the specific phenotypes associated with co-pathologies in the hippocampus and cortex. For this, we used CD68, a marker of phagocytic activity and MHCII, a key mediator of antigen presentation to distinguish activated, antigen presenting and/or phagocytic microglia. In co-pathology brains, the overall number of MHCII^+^ and MHCII^+^CD68^+^ microglia decreased compared to single pathologies (Supplemental Figure 5f,g). Conversely, MHCII^-^CD68^+^ microglia were significantly increased in the hippocampus and cortex of co-pathology mice compared to single pathology brains (Supplemental Figure 5f,g), indicating a shift towards a phagocytic response. To confirm whether these phagocytic microglia are driving the pro-inflammatory response due to co-pathologies, we assessed proliferation (Ki67 expression) and pro-inflammatory phenotype (TLR2 expression) on MHCII^+^CD68^+^ (inflammatory phagocytes) and MHCII-CD68+ (pure phagocytes) populations for functional analysis (Figure 4f). MHCII^+^CD68^+^ microglia showed decreased expression of Ki67 and TLR2 (Figure 4g,h), which could imply reduced antigen presentation in co-pathology brains. On the other hand, MHCII^-^CD68^+^ microglia showed robust proliferation and TLR2^+^ expression in the hippocampus and cortex of co-pathology brains compared to single pathology controls (Figure 4i,j). Overall, these results indicate that at 3MPI, tau, Aβ and α-syn co-pathologies synergistically promote morphological and proliferative changes in microglia, translating to a more activated, phagocytic state, likely to prioritize clearance of the increased protein pathology load seen in co-pathology brains.

### Co-pathologies promote T cell invasion and resident memory phenotype

Given the enhanced myeloid cell responses with co-pathologies (van Wetering et al., 2024) and the contribution of peripheral CD4^+^ and CD8^+^ T cells to neurodegeneration *in vivo* (Brochard et al., 2009, Ferretti et al., 2016) and in human disease (Gate et al., 2021, Haage and De Jager, 2022, Yamakawa and Rexach, 2024), we aimed to determine if T cell responses were synergistically enhanced by co-pathologies. Following co-pathology induction, brains were sectioned for IHC or dissected for single-cell isolation at 3MPI (Figure 5a). Using IHC, we show the presence of co-pathologies enhanced infiltration of CD3^+^ (Supplemental Figure 6a) or CD4+ (Figure 5b) T cells into the hippocampus compared to tau, Aβ and α-syn single pathology brains. Utilizing mononuclear cell isolation and spectral flow cytometry, our results revealed that there were significantly more CD4^+^ and CD8^+^ T cells in the hippocampus and cortex of co-pathology brains compared tau, Aβ and α-syn single pathology mice (Figure 5c-e). This data demonstrates that the presence of co-pathologies enhances the presence of both helper (CD4^+^) and cytotoxic (CD8^+^) T cells into the brain. These data suggest that adaptive immune mechanisms observed prior to neurodegeneration may be important or significantly contribute to enhanced hippocampal neurodegeneration observed with co-pathology.

**Figure 5.**
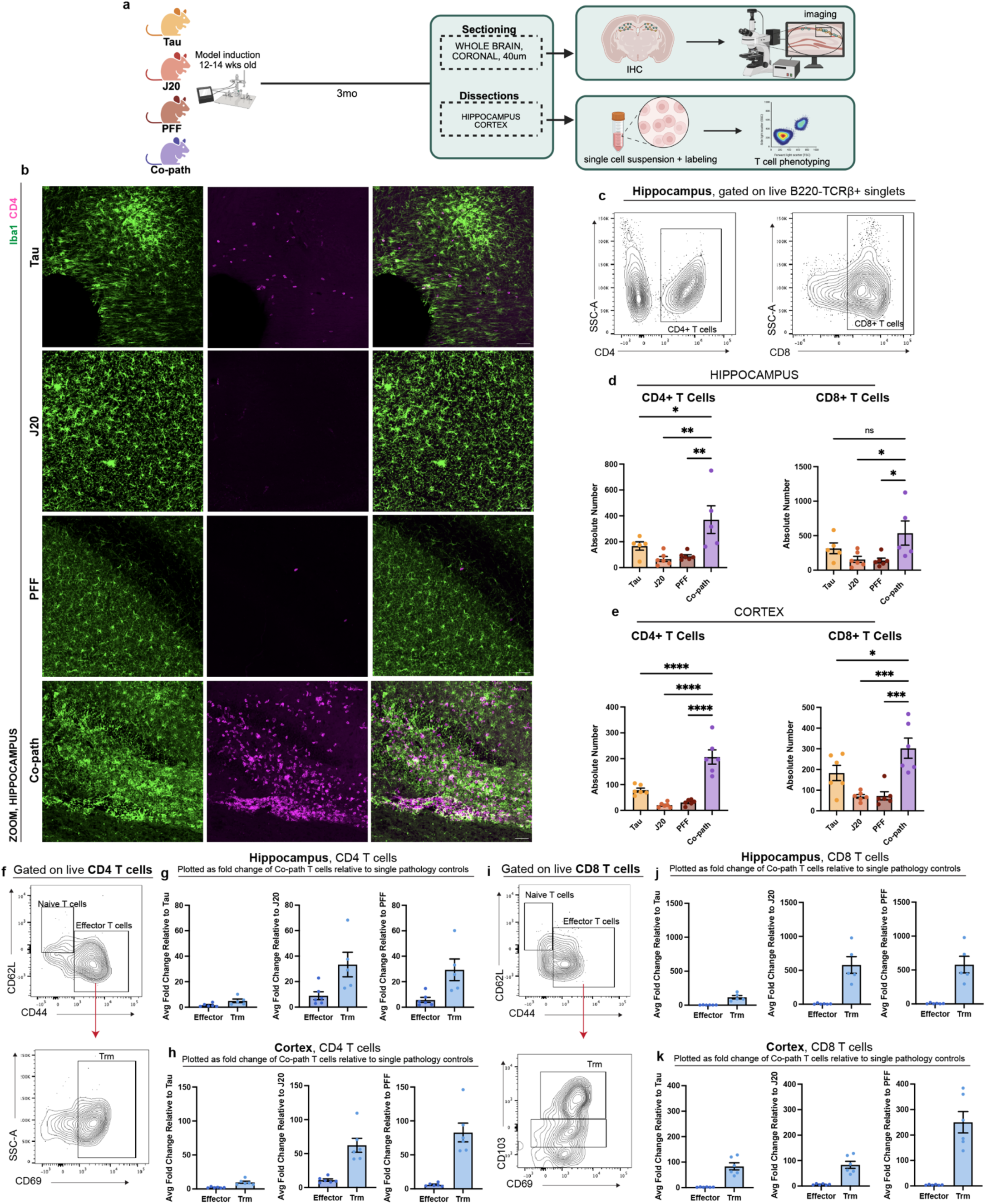
Co-pathologies promote T cell responses and tissue-resident memory T cell phenotypes at 3MPI. (a) J20 transgenic mice are injected at 12-14 weeks of age with 2uL of 5ug α-syn PFFs and 2uL of 3.2E12 vg/mL AAV9-Tau into the striatum and entorhinal cortex, respectively. For single pathology controls, age-matched non-transgenic littermates are injected with α-syn PFFs into the dorsal lateral striatum for PFF single pathology control or AAV9-Tau into the entorhinal cortex for Tau single pathology control. Age-matched J20 mice are used as the Aβ single pathology controls. At 3MPI, brains were collected and the hippocampus and surrounding cortices dissected and processed for spectral flow cytometry. (b) Representative fluorescent images of Iba1+ microglia (green) and CD4+ T cells (magenta) in the hippocampus of Tau, J20 and PFF single pathology controls compared to that in the co-pathology model. (c) Representative flow cytometry contour plots show proportion for CD4+ and CD8+ T cells. The absolute number of CD4+ and CD8+ T cells in the (d) hippocampus and (e) cortex of Tau, J20 and PFF only compared to co-pathology brains are plotted. (f) Representative contour plot showing expression and gating scheme of CD4+ Trm T cells. Quantified average fold change of CD44+ effector and CD69+ Trm in co-pathology (g) hippocampus and (h) cortex relative to Tau, J20 and PFF single pathologies. (i) Representative contour plot showing expression and gating scheme of CD8+ Trm T cells. Quantified average fold change of CD44+ effector and CD69+CD103+ Trm in co-pathology (j) hippocampus and (k) cortex relative to Tau, J20 and PFF single pathologies. Analyzed using One-way ANOVA. Mean values +/- SEM are plotted. *p<0.05, **p<0.01, ***p<0.005. n=6/group, with two brain regions pooled per sampled, both males and females included in analysis. All images taken at 10X magnification with z-stack, and digitally zoomed using Nikon Elements software. Scale bars = 50μm.

**Figure 6.**
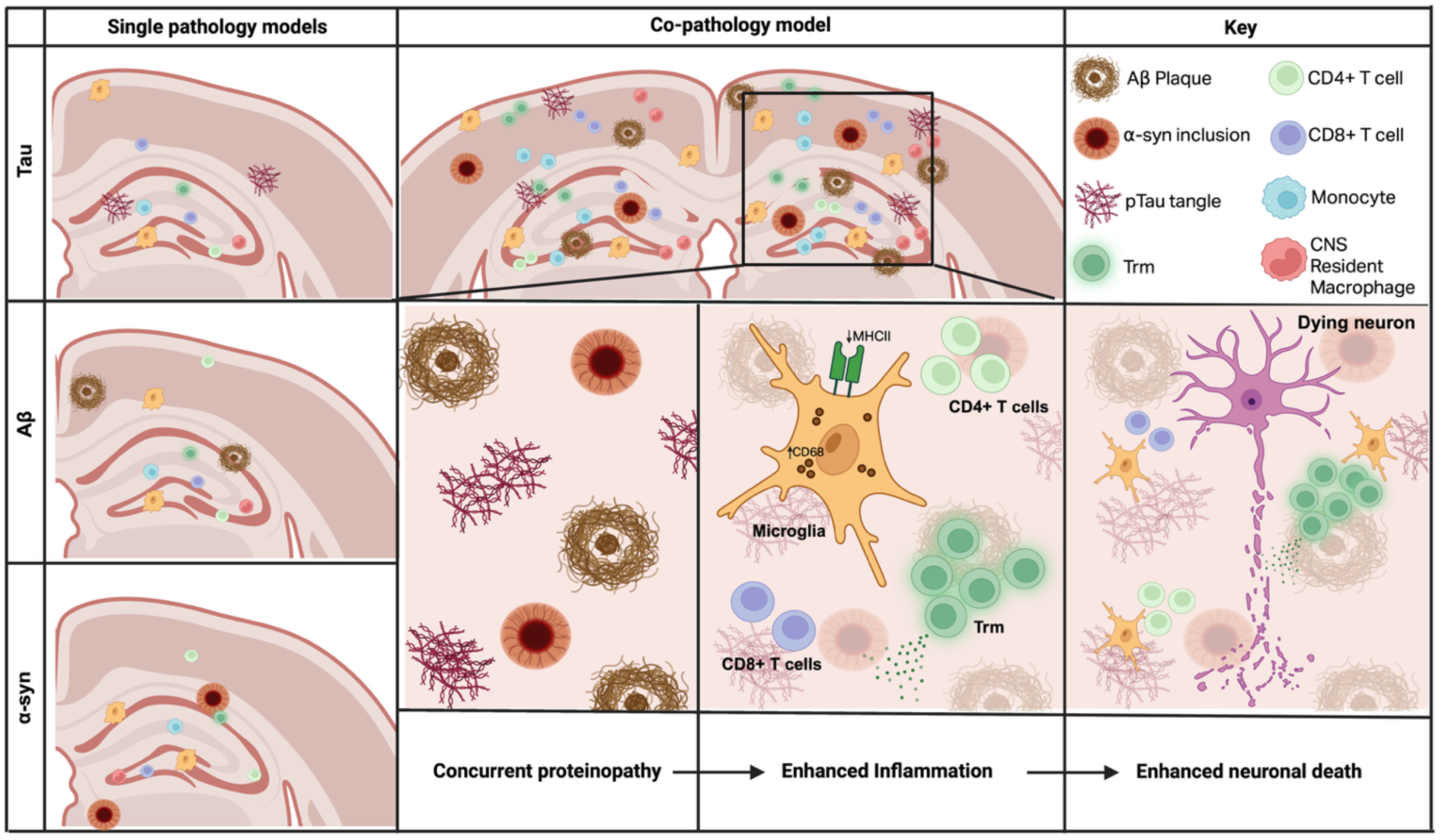
Synergistic effects of co-pathologies promote neuroinflammation, protein pathology load and neurodegeneration. Single pathology models (left panel) showing relatively mild immune activation and pathology, with little to no inclusion of tissue-resident memory (Trm) T cells, CD4+ and CD8+ T cells, monocytes and CNS resident macrophages. With the co-pathology model (middle panels), we demonstrate that concurrent protein aggregation, Aβ plaques, p-tau tangles and α-syn inclusions synergistically lead to increased recruitment and expansion of CNS-infiltrating immune cells including T cells and expanded Trm populations. This is accompanied by microglial morphology changes and elevated CD68+ expression and decreased number of MHCII+ cells. The cumulative effects of aggregated protein burden and chronic neuroinflammation result in neuronal damage and subsequent death.

Tissue resident memory (Trm) T cells have been observed in human post mortem tissue in neurodegenerative diseases such as AD and PD (Gate et al., 2020, Su et al., 2023) where they accumulate in the brain and contribute to chronic inflammation and neuronal loss (Kimura et al., 2024, Yamakawa and Rexach, 2024). Using cell surface markers expressed by Trm T cells (CD69^+^ and CD103^+^) and CD44^+^ (effector, activated T cells that are antigen experienced), we examined both CD4^+^ and CD8^+^ Trm in the hippocampus and cortex of co-pathology mice compared to single pathology mice at 3MPI. In both brain regions, co-pathologies promoted a significant expansion of both CD4^+^ and CD8^+^ Trm (CD62L^-^CD44^+^CD69^+^CD103^+^) relative to all single pathologies (Supplemental Figure 6b,c). CD44^+^CD4^+^ T cells and CD4^+^CD69^+^ Trm (Figure 5f-h) increased by 10+ fold in co-pathology brains relative to single pathology controls. CD44^+^CD8^+^ T cells and CD8^+^CD69^+^CD103^+^ Trm were increased over 100-fold in co-pathology brains compared to single pathologies (Figure 5i-k). Our findings illustrate a synergistic expansion of Trm T cells, in CD4 T cells but more robust in CD8 T cells, indicating that tau, Aβ and α-syn co-pathologies promote the long-term expansion and survival of antigen-experienced T cells within the brain prior to overt neurodegeneration.

## Discussion

Tau, Aβ and α-syn are pathological hallmarks of neurodegenerative diseases associated with cognitive decline, however, how these protein pathologies synergize to enhance neurodegeneration remains unknown. Using a novel mouse model of co-pathology incorporating tau, Aβ and α-syn co-pathologies, our results demonstrate synergistic activation of innate and adaptive immune responses in areas with co-expression of these accumulated proteins. At 3MPI, Aβ pathology and soluble alpha-synuclein levels were increased compared to the single pathology controls along with a robust neuroinflammatory response, including increased microglial cell number and changes in activation state, infiltration of T cells and enhanced Trm signature in the hippocampus and cortex. The synergistic, not merely additive, effect of co-pathologies creates a pro-inflammatory CNS environment that, in turn, enhances disease pathogenesis. Building on our previous work demonstrating that blocking neuroinflammation can prevent neuronal loss in models of neurodegeneration (Corbin-Stein et al., 2024, Williams et al., 2018), we propose that this is largely mediated through exacerbated innate and adaptive immune responses, positioning neuroinflammation as an important link between protein aggregation and neuronal loss, and a prime target for combination therapy.

Previous studies have established that multiple protein pathologies in mouse models can interact to exacerbate disease phenotypes. DLB-AD mice (3xTg AD mice crossed with α-syn A53T mice) yielded the first triple pathogenic model with worsened cognitive decline, however, there were no observed changes in microglial reactivity (Clinton et al., 2010). Combining AD-derived tau with α-syn PFFs further enhanced tau pathology and its spread relative to controls (Bassil et al., 2020). Similarly, to dissect the interaction between α-syn and Aβ, 5xFAD mice were stereotaxically injected with mouse α-syn PFFs to develop a ‘PD-AD’ mouse, that exhibited enhanced hippocampal neuronal loss, as the only mutli-pathology mouse model to assess neurodegeneration (Bassil et al., 2021). Lastly, crossing APP/PS1 mice with mutant P301L tau mice has shown increased formation of Aβ plaques and tau tangles (Lewis et al., 2001). The mechanism underlying this has been linked to Aβ plaques activating kinases, which leads to mislocalization of tau, and subsequent hyperphosphorylation (Lewis et al., 2001, Shipton et al., 2011). Results from these studies demonstrate that the presence of additional pathologies promote each other’s aggregation and accelerate cognitive decline in mice and as such, complements the work done in human postmortem studies showing co-pathology expression correlated to aging, neuroinflammation and neurodegeneration (Schneider et al., 2007, van Wetering et al., 2024). However, likely due to the design of these models, a critical question remains unanswered: whether and how tau, Aβ and α-syn synergize together as co-pathologies to trigger amplified immune responses that directly influences enhanced neurodegeneration.

Within, we created a novel co-pathology mouse model to answer this question and determine the effect of co-pathologies on pathological protein deposition, neuroinflammation, and neurodegeneration. To model this pathological overlap seen in neurodegenerative disease, we combined the ΑΑV9-double mutant Tau (tau) and α-syn mouse PFFs (α-syn) in the same J20 APP^mut^ mouse (Aβ). The J20 model provides moderate, localized Aβ pathology beginning at approximately 4 months old (Mucke et al., 2000), while AAV-driven tau propagates from the entorhinal cortex to the hippocampus and cortex to model tau spread. As endogenous mouse tau does not aggregate or form neurofibrillary tangle-like inclusions (Andorfer et al., 2003, Duff et al., 1996, Lewis et al., 2000), it was important to induce the pathology in a region like the entorhinal cortex, where localization allows for the propagation and subsequent expression of intracellular phosphorylated tau within the hippocampus and neighboring cortices. α-syn mouse PFFs induce Lewy-like inclusions with intracellular aggregation and spreading throughout the limbic and cortical regions. While AAVs that overexpress α-syn in transduced regions have been used to model PD (Decressac et al., 2012, Elabi et al., 2021, Karikari et al., 2022, Schonhoff et al., 2023), these overexpression vectors typically lack the disease-like spreading pathology and only recapitulate the molecular pathology of SNCA gene duplication/triplication carriers (Gómez-Benito et al., 2020, Pinto-Costa et al., 2023). While this model does not fully recapitulate the spatial distribution and temporal onset of pathology seen in human disease, it was designed to induce tau, Αβ and α-syn pathology within their classically affected brain regions. Like most protein overexpression and seeding models, it carries limitations, including the non-physiological levels and a lack of gradual disease progression. While postmortem tissues offer a snapshot in time of disease-driving pathology, modeling *in vivo* is needed to study interactions. Despite these caveats, this model of co-pathology offers a useful tool to investigate how these pathologies synergize to drive neurodegeneration and neuropathology.

Within, we show that the presence of co-pathologies significantly increases protein load in the brain. At 3MPI, the co-expression of tau, Aβ and α-syn in the hippocampus and cortex led to a significant increase in insoluble Aβ_1-42_ load compared to J20 single pathology brains, as well as soluble α-syn in the compared to PFF single pathology brains. These findings are consistent with previously reports showing enhanced Aβ and α-syn protein pathology load under co-pathology conditions (Bassil et al., 2020, Bassil et al., 2021). At this early timepoint, tau levels remained low and unchanged across groups in both the hippocampus and cortex, likely reflecting the slower kinetics of tau seeding and propagation mechanisms (de Calignon et al., 2012) from the entorhinal cortex to interconnected brain regions. In line with this known phenomenon, by 6MPI we observed that soluble tau was significantly increased in the hippocampus and cortex of co-pathology mice relative to Tau single pathology controls while insoluble tau levels remained comparable between groups at this later timepoint. This suggests that the early presence of Aβ pathology (already existing at time induction) together with rapidly seeded α-syn pathology creates an environment that promotes the formation of soluble tau species, including hyperphosphorylated monomers and pathogenic oligomers. Previous work has also shown that at later stages, tau burden remains similar and unchanged in a 5XFAD + αsyn multi-pathology model compared to control mice. Our co-pathology mouse model differs in that tau pathology is driven by interaction of mutant tau fragments with endogenous tau to seed rather than endogenous overexpression in the 5XFAD model, producing slower spread of pathology that manifests at the later timepoint of 6 months post-induction. In contrast to tau, we observed a decrease in overall Aβ_1-42_ and α-syn load at 6MPI, which is likely attributable to substantial loss of neurons in the hippocampus that would have produced this pathology at this stage. Collectively, these findings illustrate that co-pathologies enhance proteinopathy in the hippocampus and cortex and future combinatorial approaches may be an effective disease-modifying strategy.

Longitudinal and post-mortem studies from PD and AD cohorts show earlier onset of cognitive decline and higher mortality rates with the presence of co-pathologies (Irwin et al., 2013, Lim et al., 1999). Based on this, pure pathology is not common and may not be associated with the cognitive impairment onset in these neurodegenerative diseases (Nelson et al., 2009, Wetering et al., 2024). Here, we show that the presence of co-pathology results in enhanced neuronal loss, specifically in the CA1 and CA3, regions of the hippocampus compared to single pathology controls, correlating with region of enhanced pathology load in our model. While our dendritic spine analysis at 6MPI implied some synaptic alterations in the hippocampus, it is possible that an earlier timepoint may have captured more pronounced changes during the initial phase of synaptic disintegration, prior to this marked neuronal loss. In the SNpc, however, dopaminergic neuronal loss was prominent in the co-pathology model mice compared to the J20 mice, with no changes between co-pathology mice and Tau or PFF single pathology animals. This is likely due to the localization and abundance of the co-expression of these proteins in the hippocampus, suggesting a region-specific interaction of pathology burden. These results highlight the need for further mechanistic studies into the additive or synergistic roles of these pathologies on neurodegeneration across multiple regions.

Microglial activation and the infiltration of T cells have been documented in response to tau, Aβ, and α-syn individually in AD and PD (Chen and Yu, 2023, Hansen et al., 2018, Heneka et al., 2015, Imamura et al., 2003, Zeng et al., 2024). However, studies implicating the role of inflammation in driving neurodegeneration in a co-pathology model are lacking. Using this novel co-pathology model, we show that the co-expression of these pathologies induces robust changes in microglia, and enhanced infiltration of monocytes and T cells from the periphery indicating neuroinflammation. Compared to single pathologies, the presence of co-pathologies resulted in increased microglial cell count and activation status in the hippocampus and cortex. These changes were not limited to microglia, as we observed a significant increase in infiltrating monocyte and macrophages in areas with co-pathology burden. Our findings are consistent with recent work in human postmortem tissue that compared microglia reactivity between mixed pathology (DLB+AD) brains and pure pathology brains, showing higher levels of CD68^+^, reactive microglia in the hippocampus with mixed pathology (van Wetering et al., 2024). In our model, CD68^+^ microglia were synergistically enhanced by the presence of co-pathology and showed heightened proliferation and TLR2 expression, while MHCII^+^ microglia decreased in number without significant changes in expression of Ki67 or TLR2. Together, these findings suggest that the presence of co-pathologies shifts the microglial response towards a more proliferative, phagocytic phenotype, rather than a classic antigen-presenting role. Mechanistically, high deposition of aggregated proteins may overwhelm phagocytic pathways within microglia, diverting resources toward bulk debris clearance and away from peptide processing required for MHCII-mediated T cell interactions. This microglial phenotype is consistent with the well-established profile of disease-associated microglia (DAM), a pro-inflammatory state that is observed neurodegenerative brain diseases (Friedman et al., 2018, Keren-Shaul et al., 2017, Krasemann et al., 2017). Future work could determine whether the presence of co-pathology enhances DAM phenotypes in mice.

At 3MPI, we observed enhanced T cell responses compared to single pathologies in the hippocampus and cortex and a synergistic expansion to Trm, marked by the expression of CD69 and CD103, which are key markers for tissue retention and local immune surveillance (Smolders et al., 2018, Wakim et al., 2010). Trm have been shown to exacerbate disease through pro-inflammatory cytokine release and cytolytic functions, which may modulate the response and activation phenotypes of local microglia (Gate et al., 2020, Smolders et al., 2013). The accumulation of Trm in co-pathology brain implies their prolonged, antigen-responsive retention. Future studies should further investigate the mechanisms by which co-pathologies drive the accumulation of T cells in the CNS. Either by driving expression of tissue-residency ligands on microglia for Trm retention or alternatively by promoting compensatory antigen presentation by other CNS cells like border-associated macrophages, which have been shown promote T cell infiltration more effectively than microglia in a neurodegenerative disease model (Schonhoff et al., 2023). These findings suggest that chronic Trm-driven inflammation may represent a converging mechanism of neurodegeneration regardless of the initiating proteinopathy. As such, future depletion studies or modulation of Trm populations are necessary to determine whether co-pathology driven Trm are disease driving or potentially neuroprotective by providing a local pool of antigen experienced T cells in the CNS.

In conclusion, our findings collectively demonstrate that tau, Aβ and α-syn co-pathologies enhance proteinopathy, synergistically enhance immune activation and subsequent neurodegeneration. This synergy drives a robust immune response, shifting microglia towards a CD68+ phagocytic phenotype, and significant Trm T cells in the hippocampus and cortex. Additionally, this novel co-pathology model provides an important platform to further dissect how co-pathologies shape neuroinflammatory responses in the CNS. Ultimately, our work strongly supports the need for combinatorial therapeutic strategies that target multiple proteins and their associated immune response, moving beyond single-target approaches to effectively alter disease course.

## Methods

### Mice

The mouse strain used for this research project, B6J.Cg-*Zbtb20^Tg(PDGFB-APPSwInd)20Lms^*/Mmjax, (RRID:MMRRC_034836-JAX), was obtained from the Mutant Mouse Resource and Research Center (MMRRC) at The Jackson Laboratory, an NIH-funded strain repository, and was donated to the MMRRC by Lennart Mucke, Ph.D., Gladstone Institute of Neurological Disease. Age and sex-matched non-transgenic littermates (NTG, C57BL/6J background) were used for Sham and single pathology induction. For all experiments presented, both male and female mice were used. All research conducted on animals was approved by the Institutional Animal Care and Use Committee at the University of Alabama at Birmingham and methods followed the Animal Research: Reporting of *In vivo* Experiments (ARRIVE) 2.0 guidelines (arriveguidelines.org).

### Fibril and viral vector generation

#### Mouse α-synuclein pre-formed fibrils

α-syn mouse pre formed fibrils (PFFs) were generated and purified from mouse α-synuclein from the Volpicelli-Daley lab at the University of Alabama at Birmingham according to previously published protocols (Luk et al., 2009, Volpicelli-Daley et al., 2014). Briefly, mouse PFFs were thawed and aliquoted into 25uL (5mg/mL) and transferred to polystyrene sonication tubes. Tubes containing PFFs were then transferred to a Q700 Sonicator water reservoir once the water was at 10C. Fibrils were sonicated for 15 minutes, with pulse on and off durations, at a power of approximately 110 watts per pulse. After sonication, tubes are quickly spun down and 1uL of sonicated PFFs transferred into an Eppendorf tube in 1X PBS, for a final volume of 500uL. 5uL of this solution is sent for dynamic light scattering analysis to check for PFF fragmentation of around 50nm after sonication.

#### AAV9-Tau virus

AAV double mutant (dbl^mut^) tau (AAV9-Tau) genome plasmid was engineered by standard cloning techniques to express human tau (2N4R isoform) encoding P301L/S320F mutations driven by the hybrid chicken β-actin promoter and flanked by AAV2 iTRs.(Gray et al., 2011). Recombinant adeno-associated virus capsid 9 (AAV9) vectors were generated as described in prior studies(Sandoval et al., 2019). Briefly, HEK293 cells were co-transfected with either dbl^mut^ tau or green fluorescent protein (GFP) genome plasmids, together with plasmids encoding AAV rep/cap genes and adenovirus helper proteins. AAV viral capsids were purified from cells and culture media by iodixanol gradient centrifugation, followed by dialysis in Dulbecco’s PBS containing calcium and magnesium. Titers were determined by ddPCR using primers for the AAV iTRs and normalized to 3.2 viral genomes per mL (vg/mL).

### Stereotaxic injections

Mice aged 12-14 weeks old were deeply anesthetized using an isoflurane vaporizing instrument (UAB Animal Resource Program) and immobilized to a stereotaxic frame (Stoelting) on top of a heating pad to ensure appropriate internal body temperature is maintained. 26s/2”/2 10μL Hamilton syringes along with an automatic injecting system (Stoelting) are used to unilaterally (immunohistochemistry, DAB and fluorescence) or bilaterally (behavior, flow cytometry) inject 2uL of mouse α-syn PFFs (5μg/mL), mouse α-syn monomer (5μg/mL), AAV9-dbl^mut^ tau virus (3.2E12vg/mL) or AAV9-GFP (3.2E12vg/mL). PFFs or monomer control were injected into the dorsal lateral striatum based on bregma (AP +0.5mm, ML +/-2.0mm and DV -3.0mm from the dura) AAV9-Tau or AAV9-eGFP viruses were injected into the entorhinal cortex (AP -3.5mm, ML +/-3.9mm and DV -4.2mm from dura). Injection needle was left in the injection site for an additional 2 minutes following completion and then slowly retracted from the site over the course of another 2 minutes. All surgical protocols and post-surgery aftercare methods were followed and approved by the Institutional Animal Care and Use Committee at the University of Alabama at Birmingham.

To induce the co-pathology model, 3-month-old J20 transgenic mice were injected with mouse α-syn PFFs into the striatum and AAV9-Tau virus into the entorhinal cortex. For sham surgery, non-transgenic (NTG) littermates were injected with mouse α-syn monomer and AAV9-eGFP into the striatum and entorhinal cortex, respectively.

To induce single pathology controls, NTG mice were injected with mouse α-syn PFFs into the dorsal lateral striatum for α-syn models and AAV9-Tau virus into the entorhinal cortex for tau models. α-syn monomer into the striatum and AAV9-GFP were injected into the dorsal lateral striatum and entorhinal cortex, respectively, of non-transgenic mice as controls. J20 transgenic mice were used as the Aβ single pathology mice, compared to un-injected NTG littermates.

### Tissue preparation and processing

At 3 months post-induction (3MPI) and 6 months post-induction (6MPI) of the co-pathology model and single-color controls, mice were anesthetized using isoflurane and transcardially perfused with 0.01M phosphate-buffered saline (PBS, pH = 7.4) and fixed with 4% paraformaldehyde (PFA in PBS, pH = 7.4). Brains were then dissected and incubated in 4% PFA for 2 hours at room temperature. After fixation, brains were placed in 30% sucrose in PBS for 3 days until brains were fully saturated. Brains were then frozen on dry ice and cryosectioned coronally at 40μm on a sliding microtome (Thermo Scientific Micron HM 450). Sliced brain tissue was then stored in a 50% glycerol/PBS solution at -20°C.

### Immunofluorescent immunohistochemistry

Free-floating 40-μm thick sections were washed in 0.01M tris-buffered saline solution (TBS, pH 7.4) prior to an antigen retrieval process with a solution of 10mM sodium citrate and 0.05% Tween-20 in TBS. Sections were then incubated in 5% blocking serum at room temperature before being incubated overnight in the following primary antibody solutions (TBST, 1% serum): anti-CD4 (clone RM4-5, ThermoFisher), anti-CD8 (clone 4SM15, eBioscience) and anti-Iba1 (WACO). On the following day, sections were washed in TBS with .1% Triton-100 and incubated in fluorescent conjugated antibody in the dark for 2 hours at room temperature. Sections were then washed in TBST and mounted on microscope slides for analysis. Slides were imaged at 20X on a Nikon A1 Laser Scanning Confocal microscope.

### DAB immunohistochemistry

Free-floating 40-μm thick sections were washed in 0.01M tris-buffered saline solution (TBS, pH 7.4) prior to being quenched with 3% hydrogen peroxide in TBS for 5 minutes. Next, sections underwent an antigen retrieval process with a solution of 10mM sodium citrate and 0.05% Tween-20 in TBS for 30 minutes at 37°C. Following this, sections were blocked in a 5% goat or donkey serum solution for at least 1 hour at room temperature and then incubated overnight at 4°C with the following antibodies in 1% serum solution: anti-CD3 (clone 17A2, eBioscience), anti-tyrosine hydroxylase (clone: EP1536Y, Abcam), anti-alpha synuclein, phosphor-serine 129 (pSer129, close EP1536Y, Abcam), anti-AT8 (clone MN1020, ThermoFisher), anti-Aβ (clone D54D2, Cell Signaling) and anti-NeuN (clone EPR12763, Abcam). Sections were then incubated with a biotinylated secondary antibody (1:1000, Vector Labs) in a 1% serum in TBST solution. After incubation, sections were washed with TBS and incubated in the Vectastain ABC Reagent (Vector Labs) for 30 minutes at RT prior to development with the Vectastain diaminobenzidine DAB Kit (SK-4199, Vector Labs) according to the manufacturer’s protocol. Sections were then mounted onto plus-coated slides and dehydrated with a gradient of ethanol solutions (70%-100%) followed by Citrisolv. Slides were them coverslipped with Permount Mounting Media (Electron Microscopy Sciences). Slides were then images at 10X and 20X on a Zeiss Imager M2 brightfield microscope (MBF Biosciences).

### Unbiased stereology of the hippocampus and substantia nigra pars compacta

In order to quantify neuronal cell loss, an unbiased stereology protocol was used. TH and NeuN-DAB stained (see ‘DAB Immunohistochemistry’ section) SNpc and hippocampal tissue sections, respectively, were imaged and analyzed with a Zeiss Axio Imager M2 microscope (MBF Biosciences) and the Cavalieri stereology probe used to count cells, with a section thickness of 25μm to account for dehydration of sections. To quantify NeuN+ neurons in the hippocampus, contours were drawn around the five to seven sections, focusing on the CA1 and CA3 regions. To quantify Th+ neurons in the SNpc, contours were drawn around both sides of the nigra, in four to five sections. Similar contours were drawn in the nigra to quantify α-syn+ neurons. For stereological counting, the grid size used for the hippocampus was 130μmx130μm and the counting frame was 40μmx40μm. In the SNpc, the grid size and counting frame were, 100x100 and 50μmx50μm, respectively.

### Brain single cell isolation and flow cytometry

At 3MPI, co-pathology and single pathology mice were anesthetized and transcardially perfused with 0.01M PBS (pH 7.4). Briefly, brain tissue was removed and the brain regions dissected. Tissue was then digested with 1 mg/mL Collagenase IV (Sigma) diluted in RPMI 1640 (1X, Gibco). Following enzyme digestion, tissue was then filtered and smashed through a 70μm filter using a syringe plunger and mononuclear cells isolated out using a 30/70% Percoll gradient (GE).

For cell staining, isolated brain cells were blocked with anti-Fcγ receptor (1:100, Biosciences). Following this, cell surface markers were labelled with the following, fluorescent-conjugated antibodies against CD45 (clone 30-F11, eBioscience), CD11b (clone M1/70, BioLegend), MHCII (M5/114.15.2, BioLegend), Ly6C (clone HK 1.4, BioLegend), TCRβ (clone H57-597, Biolegend), CD4 (GK1.5, BioLegend), CD8a (clone 53-6.7, BioLegend). CD69 (clone H1.2F3, BioLegend), B220 (clone RA3-6B2, BD Biosciences), CD44 (clone IM7, Biolegend), CD62L (clone MEL-14, BioLegend), CD103 (clone 2E7, BioLegend), TLR2 (clone CB225, BD Biosciences), Ki67 (clone 16A8, BioLegend), CX3CR1 (clone SA011F11, BioLegend), CD68 (clone FA-11, BioLegend). A fixable viability dye was used to distinguish live cells, per the manufacturer’s protocol Aqua-LIVE/DEAD Stain Kit (Invitrogen). As single-color compensation controls, isolated splenocytes or UltraComp eBeads^TM^ (Invitrogen) were used. To analyze and run samples, a BD Symphony (BD Biosciences) or Spectral Enabled Symphony flow cytometer were used at the UAB Flow Cytometry and Single Cell Core Facility. Mean cell counts, percentages and mean fluorescent intensity were measured and analyzed using FlowJo (Tree Star) software.

### Mouse brain lysate preparation

Brain tissue was micro dissected, snap frozen in super-cooled bath of 2-methylbutane with dry ice, and stored at -80°C. Tissue lysates were homogenized with a tissue grinder using 20% weight/volume lysis buffer (50 mmol/L Tris-HCl pH 7.4, 175 mmol/L NaCl, 5 mmol/L EDTA) supplemented with protease and phosphatase inhibitors and sonicated for 10 sec on ice. After addition of Triton X-100 to a final concentration of 1% volume/volume, samples were incubated on ice for 30 minutes and centrifuged at 15,000 xg for 60 min at 4°C. The supernatant was saved as the Triton X-100 soluble fraction. The pellet was resuspended in lysis buffer with 2% SDS, sonicated for 10 sec, and centrifuged for 10 minutes at 15,000 xg. The resulting supernatant was reserved as the Triton X-100 insoluble fraction. Protein concentration was determined by the bicinchoninic acid assay (Pierce Cat# 23225).

### Amyloid β detection by Aβ1-42 ELISA

A total of 20 µg brain lysate was used to measure concentration of Aβ1-42 with the human Amyloid beta (aa1-42) Quantikine ELISA (R&D Systems Cat# DAB142) kit following the manufacturer’s protocol. Colorimetric signal was measured using the SpectraMax iD3 (Molecular Devices).

### Western Blot

#### Chemiluminescence

A total of 20µg of protein from the Triton X-100 soluble and insoluble fractions described above were denatured in dithiothreitol (DTT) buffer (0.25 M Tris-HCl pH 6.8, 200mM DTT, 8% SDS, 30% glycerol, 2mg bromophenol blue) by boiling for 10m. Samples were run on 15% SDS-polyacrylamide gels and transferred onto nitrocellulose membranes, which were then fixed for 30m in 0.4% paraformaldehyde. Blots were blocked in 5% dry milk in TBS-T for 1 hour at ambient temperatures and incubated in primary antibodies at 4°C overnight. Primary antibody concentrations are as follows: Alpha-Synuclein (1:1000; BD-Biosciences 610787) and GAPDH (1:10,000; Cell Signaling 14C10). Blots were washed in TBS-T, incubated in HRP-conjugated secondary antibodies (Mouse HRP: Jackson Labs 115-035-003; Rabbit HRP: Jackson Labs 111-035-003) at 1:2000 for 1h at ambient temperatures, and imaged using a BioRad ChemiDoc MP system. Blots were analyzed using BioRad Image Lab software.

#### Fluorescence

Blots were run as described above. LI-COR blocking buffer and antibody diluents (LI-COR 927-66003) were used with Tau (1:1000; Cell Signaling 43984S) and GAPDH antibodies. IRDye-conjugated secondary antibodies (Mouse IRDye 680: LI-COR 926-68070; Rabbit IRDye 800: LI-COR 926-32211) were used at 1:2000 for 1h at ambient temperatures. Blots were imaged using a LI-COR Odyssey CLx system and analyzed using LI-COR Image studio software.

### Dendritic spine morphology analysis

#### Perfusions and brain tissue processing

For iontophoretic microinjections, mice were anesthetized with Fatal Plus (Vortech Pharmaceuticals, Catalog #0298-9373-68) and transcardially perfused with cold 1% paraformaldehyde (PFA; Sigma Aldrich, Catalog #P6148) for 1 min, followed by 4% PFA with 0.125% glutaraldehyde (Fisher Scientific, Catalog #BP2547) for 10 min. A peristaltic pump (Cole Parmer) was used for consistent administration of cold PFA. Immediately following perfusion, each brain was extracted and drop-fixed in 4% PFA with 0.125% glutaraldehyde for 8–12 h at 4 °C. After fixation, brains were coronally sliced in 250 µm thick sections using a Leica vibratome (VT1000S, speed 70, frequency 7). Brains were sliced in 0.1 M PB and stored at 4 °C, one slice per well in a 48-well plate, in 0.1 M PB with 0.1% sodium azide (Fisher, Catalog #BP922I).

#### Iontophoretic microinjections

Iontophoretic microinjections were performed based on methods described previously(Greathouse et al., 2019, Henderson et al., 2019). A Nikon Eclipse FN1 upright microscope with a 10× objective and a 40× water objective was placed on an air table. The tissue chamber used consisted of a 50 × 75 mm plastic base with a 60 × 10 mm petri dish epoxied to the base. A platinum wire was attached so that the ground wire could be connected to the bath by an alligator clip. The negative terminal of the electric current source was connected to a glass micropipette filled with 2 µL of 8% Lucifer yellow dye (ThermoFisher, Catalog#L453). Micropipettes (A-M Systems, Catalog #603500) with highly tapered tips were pulled fresh at the time of use. A manual micromanipulator was secured on the air table with magnets that provided a 45° angle for injection. Brain slices were placed into a small petri dish containing 1× PBS and DAPI for 5 min at room temperature. After incubation in DAPI, slices were placed on dental wax, and then a piece of filter paper was used to adhere the tissue. The filter paper was then transferred to the tissue chamber filled with 1× PBS, and weighted down for stability. The 10× objective was used to visualize advancement of the tip of the micropipette in X, Y, and Z planes until the tip was just a few micrometers above the tissue. The 40× objective was then used while advancing the tip into layer 2/3 of the ventral CA1 regions of the hippocampus. Once the microelectrode contacted a neuron, 2 nA of negative current were applied for 5 min to fill the neuron with Lucifer yellow. After 5 min, the current was turned off and the micropipette was removed from the neuron. Neuron impalement within the CA1 occurred randomly in a blind manner. If the entire neuron did not fill with dye after penetration, the electrode was removed and the neuron was not used for analysis. Multiple neurons were injected in each hemisphere of each animal. After injection, the filter paper containing the tissue was moved back into the chamber containing 1× PBS. The tissue was carefully lifted off the paper and placed on a glass slide with two 125 µm spacers (Electron Microscopy Sciences, Catalog #70327-20S). Excess PBS was carefully removed with a Kimwipe, and the tissue was air-dried for 1 min. One drop of Vectashield (Vector Labs, Catalog #H1000) was added directly to the slice, and the coverslip (Warner, Catalog #64-0716) was added and sealed with nail polish. Injected tissue was stored at 4 °C in the dark.

#### Confocal microscopy

Confocal microscopy was used to capture images of Lucifer yellow-filled dendrites on pyramidal neurons in CA1 of the hippocampus. Our methods are based on previously described methods(Greathouse et al., 2018, Henderson et al., 2019), and are detailed as follows. Imaging was performed by a blinded experimenter on a Nikon (Tokyo, Japan) Ti2 C2 confocal microscope, using a Plan Apo 60×/1.40 NA oil-immersion objective. Three-dimensional z-stacks were obtained of secondary dendrites from dye-impregnated neurons that met the following criteria: (1) within 80 µm working distance of microscope, (2) relatively parallel with the surface of the coronal section, (3) no overlap with other branches, (4) located between 40 and 120 µm from the soma. Nikon Elements 4.20.02 image capture software was used to acquire z-stacks with a step size of 0.1 µm, image size of 1024 × 512 px, zoom of 4.8×, line averaging of 4×, and acquisition rate of 1 frame/s.

Confocal microscopy was also used to capture images of slices from mice. Imaging was performed on a Nikon Ti2 C2 confocal microscope, using a Plan Fluor 10×/0.75 NA air objective. Nikon Elements 4.20.02 image capture software was used to acquire stitched images (20 × 10 field) by automatically stitching multiple adjacent 10× images using 25% blending overlap. Each frame had an image size of 1024 × 1024 px, and acquisition rate of 1 frame/s. Subsequently, 10× stitched images were acquired, with all other imaging parameters remaining the same. Following image acquisition, post-image processing was performed using Fiji ImageJ. Each image file was a two channel image. The Split Channels function was used to separate the two channels.

#### Dendritic spine morphometry analysis

Confocal z-stacks of dendrites were deconvolved using Huygens Deconvolution System (16.05, Scientific Volume Imaging, the Netherlands). The following settings were used: deconvolution algorithm: GMLE; maximum iterations: 10; signal to noise ratio: 15; quality: 0.003. Deconvolved images were saved in .tif format. Deconvolved image stacks were imported into Neurolucida 360 (2.70.1, MBF Biosciences, Williston, Vermont) for dendritic spine analysis. For each image, the semi-automatic directional kernel algorithm was used to trace the dendrite. The outer 5 µm of each dendrite were excluded from the trace. All assigned points were examined to ensure they matched the dendrite diameter and position in X, Y, and Z planes, and were adjusted if necessary. Dendritic spine reconstruction was performed automatically using a voxel-clustering algorithm, with the following parameters: outer range = 5 µm, minimum height = 0.3 µm, detector sensitivity = 80%, minimum count = 8 voxels. The semi-automatically identified spines were examined to ensure that all identified spines were real, and that all existing spines had been identified. If necessary, spines were added by increasing the detector sensitivity. Additionally, merge and slice tools were used to correct errors made in the morphology and backbone points of each spine. Each dendritic spine was automatically classified as a dendritic filopodium, thin spine, stubby spine, or mushroom spine based on constant parameters(Greathouse et al., 2019, Henderson et al., 2019). Three-dimensional dendrite reconstructions were exported to Neurolucida Explorer (2.70.1, MBF Biosciences, Williston, Vermont) for branched structure analysis, which provides measurements of dendrite length; number of spines; spine length; number of thin, stubby, and mushroom spines, and filopodia; spine head diameter; and spine neck diameter, among other measurements. The output was exported to Microsoft Excel (Redmond, WA). Spine density was calculated as the number of spines per 10 µm of dendrite length. Values per mouse were calculated by averaging the values for all dendrites corresponding to that mouse. To assess apical and basal dendrites separately, values for either apical or basal dendrites were averaged per mouse. For a mouse to be included in statistical analyses for dendritic spine density and morphology a minimum of 100 µm of dendrite length had to be reconstructed and analyzed.

### Behavior

Behavior tests to assess for motor and cognitive function were conducted at 6 months post-induction of co-pathology model or single pathology models. Mice were handled for 3 days prior to the start of testing and were habituated to the testing room for 1 hour at the start of each testing day. The order of test was designed to minimize stress to the animals, such that lower-stress tasks were completed before high-stress tasks. All apparatus was cleaned with 2% chlorohexadine between trials. The researcher conducting the tests was blinded to experimental groups during trials and analysis.

#### Open Field

To test for general motor activity, mice were place in a 30.48 x 30.48 cm enclosed field with opaque walls and allowed to explore for 10 minutes. The inner zone was defined as the center of the box. Noldus Average velocity, distance traveled, and percent of the time spent in the center were determined by the Ethovision XT v16 software.

#### Pole Test

To further test for motor deficits, mice were placed in a dark room and habituated for 1 hour starting at the beginning of the dark phase. Two red lamps were strategically placed in the room to visualize the behavior test without disrupting the dark phase. The pole test was performed using a 53cm wooden dowel rod with a diameter of 10 mm, anchored with a 13.9 x 20 x 1.8 cm block of wood and ridged at 2.54 cm intervals with rolled rubber bands. The dowel rod was placed vertically in the center of a 1x1 foot enclosure. The mice were first trained by being placed face-down at the base of the pole and allowed to climb off onto the floor of the enclosure before being returned to their cage. This training was done twice. Following this, each mouse was placed face-up at the top of the pole and allowed no more than 180 seconds to turn around and climb down to the bottom; this was repeated five times. Mice were given a minimum of 30 seconds to rest between each trial. Three time point measurements were recorded. The first time point started from the time the mouse was placed on the pole to the time taken for the mouse to reach the bottom (total descent time). Measurements were recorded for the time the mouse took to turn around on the pole to face the bottom (turnaround time) and the total time to descend (time descending) in seconds. The average time for each time point was calculated from all five trials for each mouse.

#### Rotarod

To further test for motor activity, mice were placed on an accelerating rotarod apparatus, and time taken to fall was measured. Mice were trained on the apparatus over 5 days, with the rotarod speed increasing from 4 rpm to 40 rpm over 300 seconds. If mice had not fallen by 120 seconds during the testing trial, the rotation was paused, mouse was removed, and the time was recorded as 120 seconds. Mice underwent two trials per day with 30 minutes of resting time between each trial.

#### Barnes Maze

A blue circular apparatus of 122cm in diameter consisting of forty, five cm, evenly spaced holes was used throughout the test. The test consisted of three stages: (1) Habituation phase, (2) Acquisition phase, and (3) Probe phase. The habituation phase took place on day 1 where mice were placed in the center of the apparatus covered by a black mesh cup uniformly covered. 10 seconds after the placement, mice freely roamed the Barnes maze for an undefined amount of time until they found and independently entered the escape tunnel. The acquisition phase occurred through Days 2-6 and consisted of one, five-minute trial. If mice did not find nor enter the escape tunnel after the five-minute trial, mice were gently guided towards the direction of the tunnel where they then entered it on their own. Entry into the tunnel was defined as all four paws in the tunnel. Upon entry, a cover was used to cover to hole to prevent the mice from leaving the tunnel. The tunnel was then carefully removed from the maze apparatus and gently placed in the home cage of the mouse. The latency to find and enter the escape tunnel was recorded. On Day 10, mice were again introduced to the maze for one minute without the escape tunnel present. The amount of time spent in the escape tunnel quadrant, entry zone, and goal box were determined by the Ethovision XT v16 software.

### Statistical Analysis

All graphs and subsequent statistical tests were generated using Prism software (GraphPad). Flow cytometry experiments used ten to twelve independent samples with two brain regions pooled per sample, for a final number of five to six per group for experiments. Data were analyzed using an unpaired student’s t-test or a one-way ANOVA. Graphs within display the mean +/- SEM. For unbiased stereology, estimated populations were generated and a one-way ANOVA was used to determine the significance between groups. *p<0.05, **p<0.01, ***p<0.0005, ****p<0.0001.

### Data availability

The authors affirm that the findings within the manuscript are supported by the data therein. The data, protocols and key lab materials used and generated in this study are listed in a Key Resource Table alongside their persistent identifiers on Zenodo (10.5281/zenodo.16739907).

## Declarations

## Acknowledgements

We would like to thank Josh Randolph for maintenance and assistance with animal husbandry and handling. We thank Rajesh Gupta for his preliminary contributions to immunohistochemistry staining. We thank Veronica Obregon-Perko for her assistance with spectral flow cytometry panel design. We also thank Laura Volpicelli-Daly, Charlotte Brzozowski and Marissa Menard for their contribution of mouse α-syn PFFs for our studies. Lastly, we would like to thank Warren Hirst, Lucas Hampton and Karen Eskow-Jaunarajs for their input and guidance during the development of this project. We thank the UAB Comprehensive Flow Cytometry Core for their assistance with flow cytometry experiments. The core is supported by the Center for AIDS Research, AI027767, The O’Neal Comprehensive Cancer Center, CA013148 and Shared instrument grant S10OD032296. Experimental design figures were generated using BioRender. This work was supported in whole by Aligning Science Across Parkinson’s (Grant# ASUB00000984) through the Michael J. Fox Foundation for Parkinson’s Research (MJFF). For the purposes of open access, the author has applied a CC BY public copyright license to all Author Accepted Manuscripts arising from this submission.

## Author’s contributions

JMW and ASH designed the study, experiments, and wrote the manuscript. JMW was responsible for planning and performing all stereotaxic surgeries. JMW, YTY, ATM, NJCS, WJW and AZ contributed towards tissue collection and executing flow cytometry experiments. TAY, WJS, KS and JMW performed all ELISA and immunoblotting experiments and analysis. JMW, NM and ACS conducted IHC stains. JMW manually assessed NeuN+ neuronal loss via unbiased stereology. AZ manually assessed Th+ neuronal loss via stereology. NHC, LFL, PNM, KMG and JHH planned, conducted and analyzed dendritic spine morphology counts. FPM and IS developed and optimized the AAV double-mutant tau vector. JMW and ASH drafted and constructed the manuscript and figures. FPM, JHK and DJT provided feedback on study design and edits to the manuscript. All authors read and approved the final manuscript.

## Supplemental Figures

**Supplemental Figure 1.**
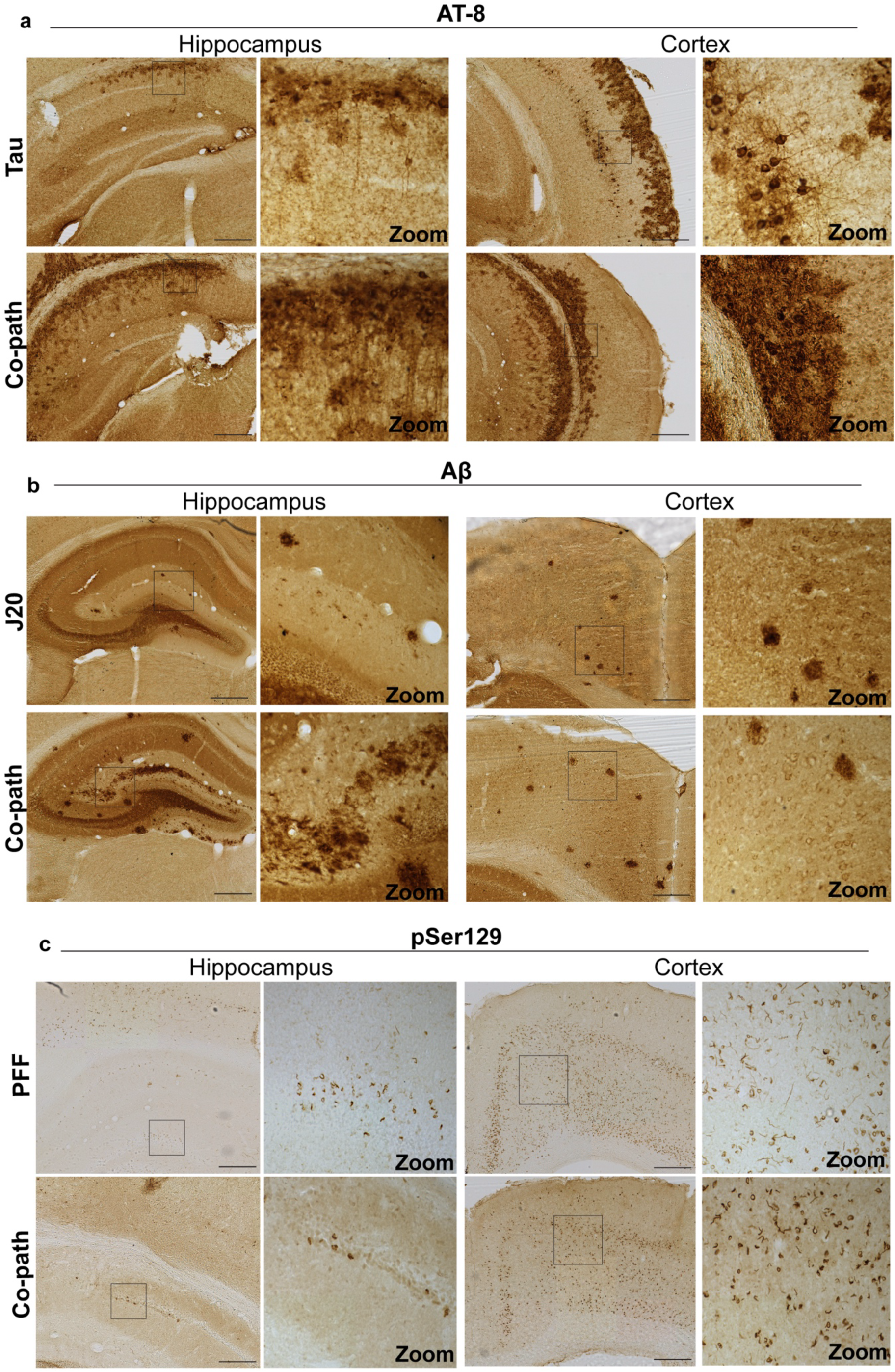
AT-8, Aβ or pSer129 protein pathology at 6MPI in the hippocampus and cortex of co-pathology and single pathology brains. (a) AT-8/pTau-positive neurons (DAB+, brown) in the hippocampus and cortex of tau single pathology and co-pathology brains. (b) Aβ-positive plaques (DAB+, brown) expressed in the hippocampus and cortex of J20 single pathology and co-pathology brains. (c) pSer120-positive inclusions in the hippocampus and cortex of PFF only (α-syn) and co-pathology brains. All images taken at 10X or 20X for zooms, scale bars = 100μm.

**Supplemental Figure 2.**
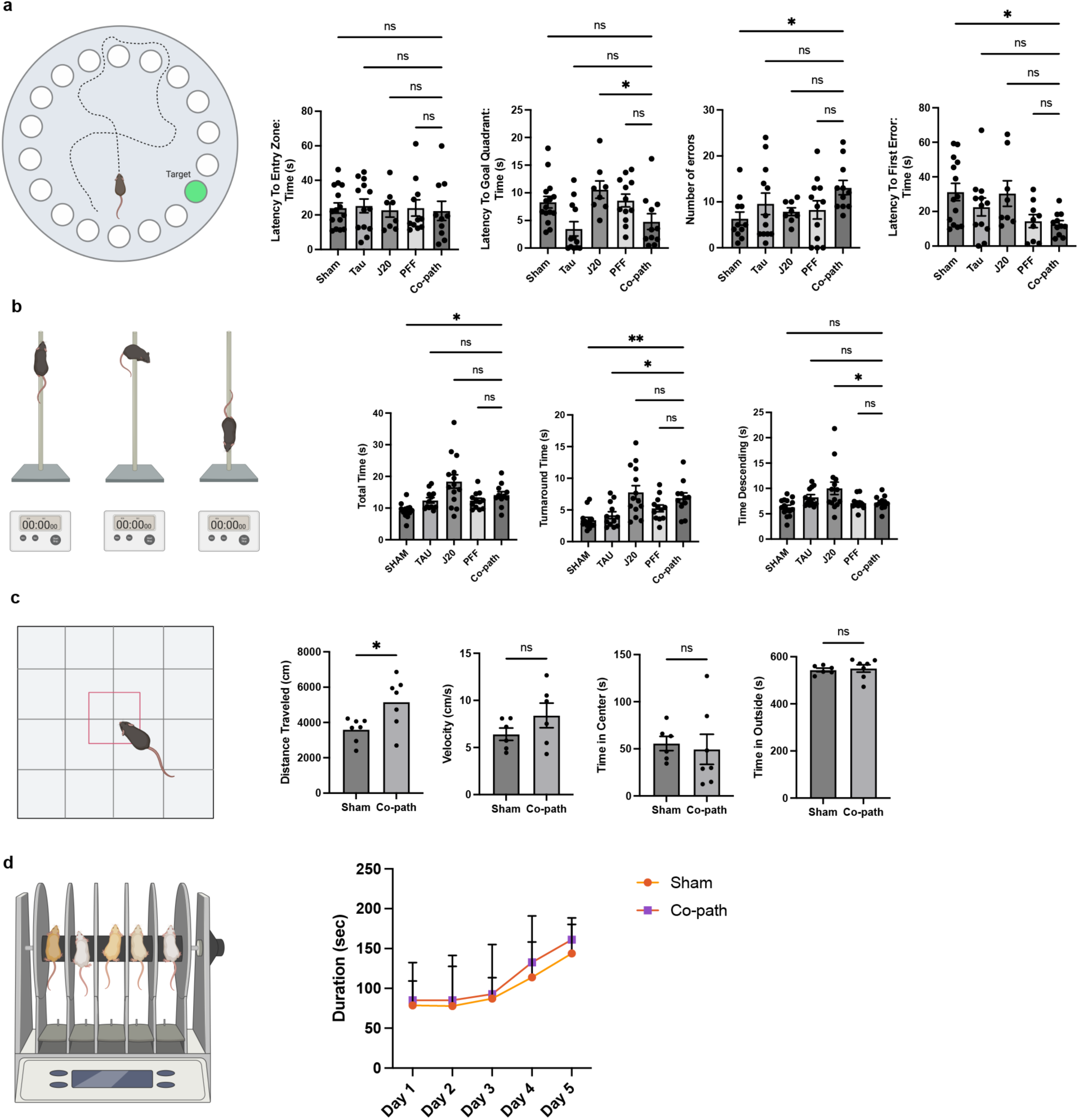
Motor and cognitive behavioral tests to assess for impairments in co-pathology vs single pathology animals. (a) Quantification of latency to entry zone, to goal quadrant, to first error and the total number of errors analyzed from Barnes maze test. (b) Measures of total time, turnaround time and time descending on the pole test. (c) Distance travelled, velocity, time in center and time in outside are depicted from open field test. (d) Total duration of animals on rotarod platform across 5 days of testing. Student’s t-test or One-way ANOVA with post hoc for significance. Mean values +/- SEM are plotted. ns = no significance, *p<0.05, **p<0.01, ***p<0.005. n=8-10 animals per group, both males and females used.

**Supplemental Figure 3.**
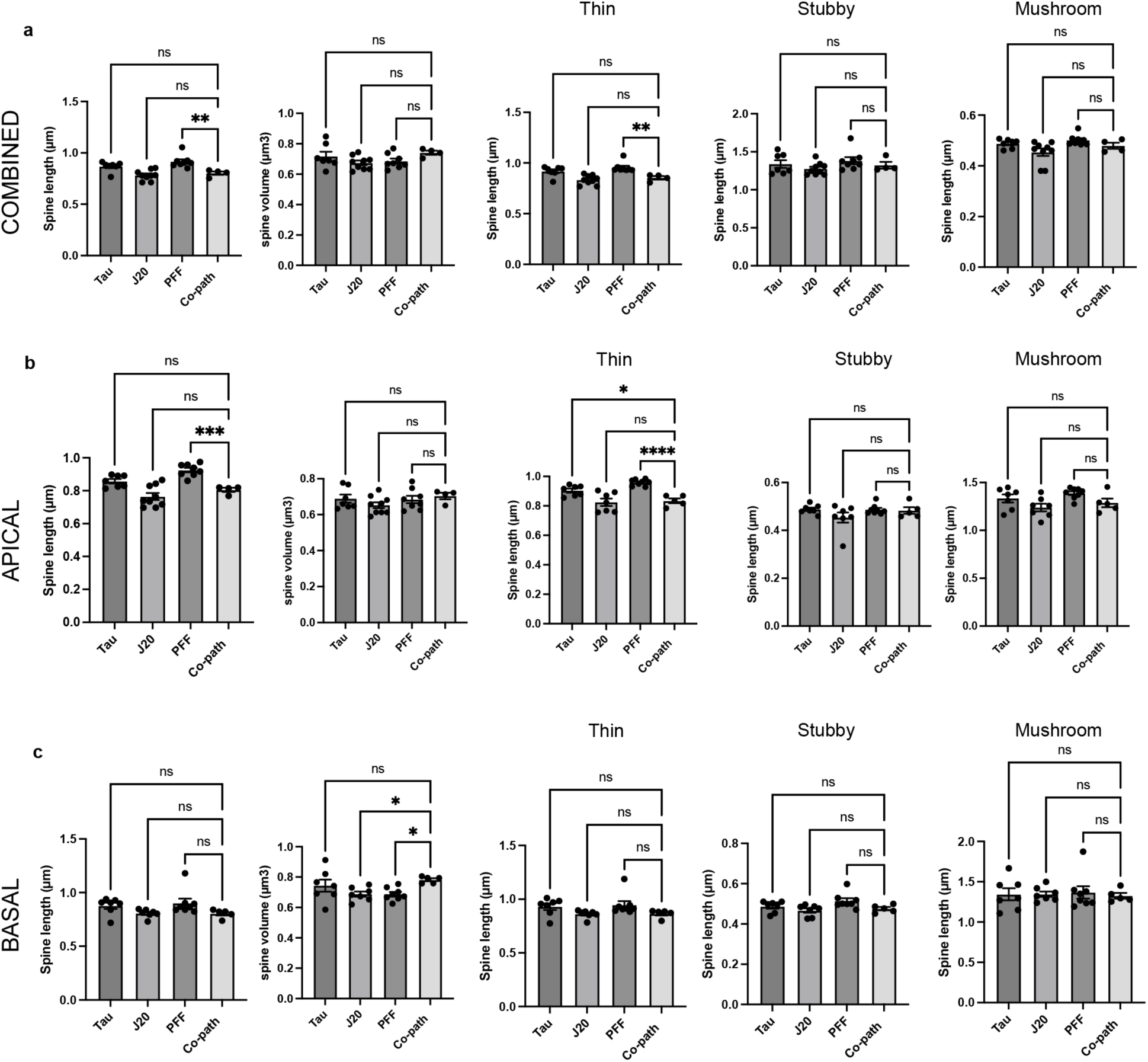
Assessment of synapse integrity in the hippocampus of co-pathology and single pathology mice at 6MPI. Dendritic spine morphology was assessed in the CA1 region of hippocampus on pyramidal neurons and measures of total spine length and spine volume for all spines and for thin, stubby and mushroom subgroups are plotted from (a) combined (both apical and basal), (b) apical only or (c) basal only dendrites. One-way ANOVA with post hoc for significance. Mean values +/- SEM are plotted. ns = no significance, *p<0.05, **p<0.01, ***p<0.005. n=8-10 animals per group, both males and females used.

**Supplemental Figure 4.**
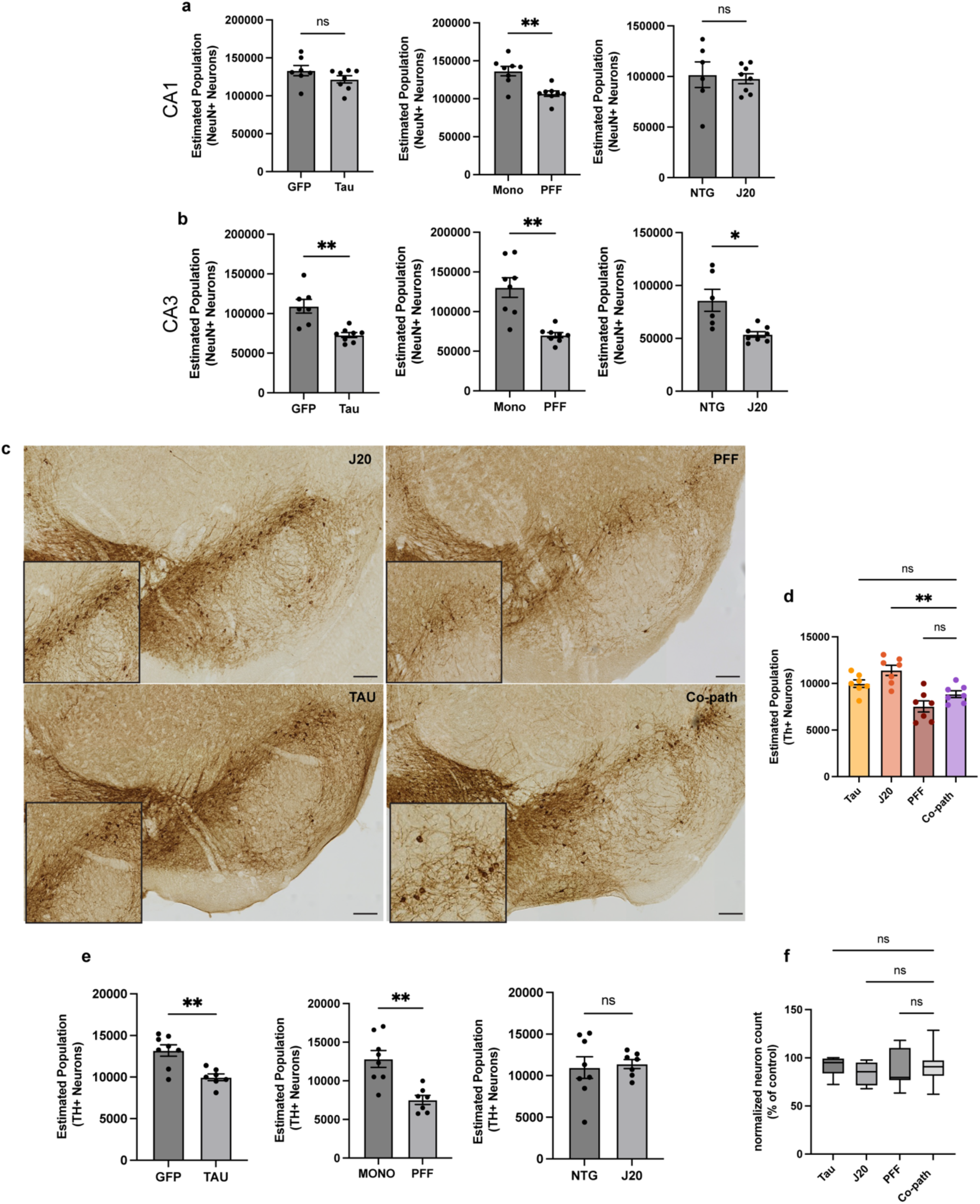
Unbiased stereology for quantification of NeuN+ neurons in the hippocampus (single pathology and controls) and Tyrosine Hydroxylase positive neurons in the substantia nigra pars compacta (SNpc) at 6MPI. (a,b) Estimated population of NeuN+ neurons of co-pathology and single pathology CA1 or CA3 compared to their individual controls. (c) TH+ neurons (brown) in the SNpc of single pathology and co-pathology brains. (d) Unbiased stereology was used to quantify the number of TH+ neurons in the SNpc in the co-pathology mouse model compared to Tau, J20 and PFF single pathology brains. All images taken at 20X. Scale bars = 100μm. Student’s t-test or One-way ANOVA with post hoc for significance. Mean values +/-SEM are plotted. Ns = no significance, **p<0.01, ***p<0.005, ****p<0.0001. n=7-8 animals per group, both males and females used.

**Supplemental Figure 5.**
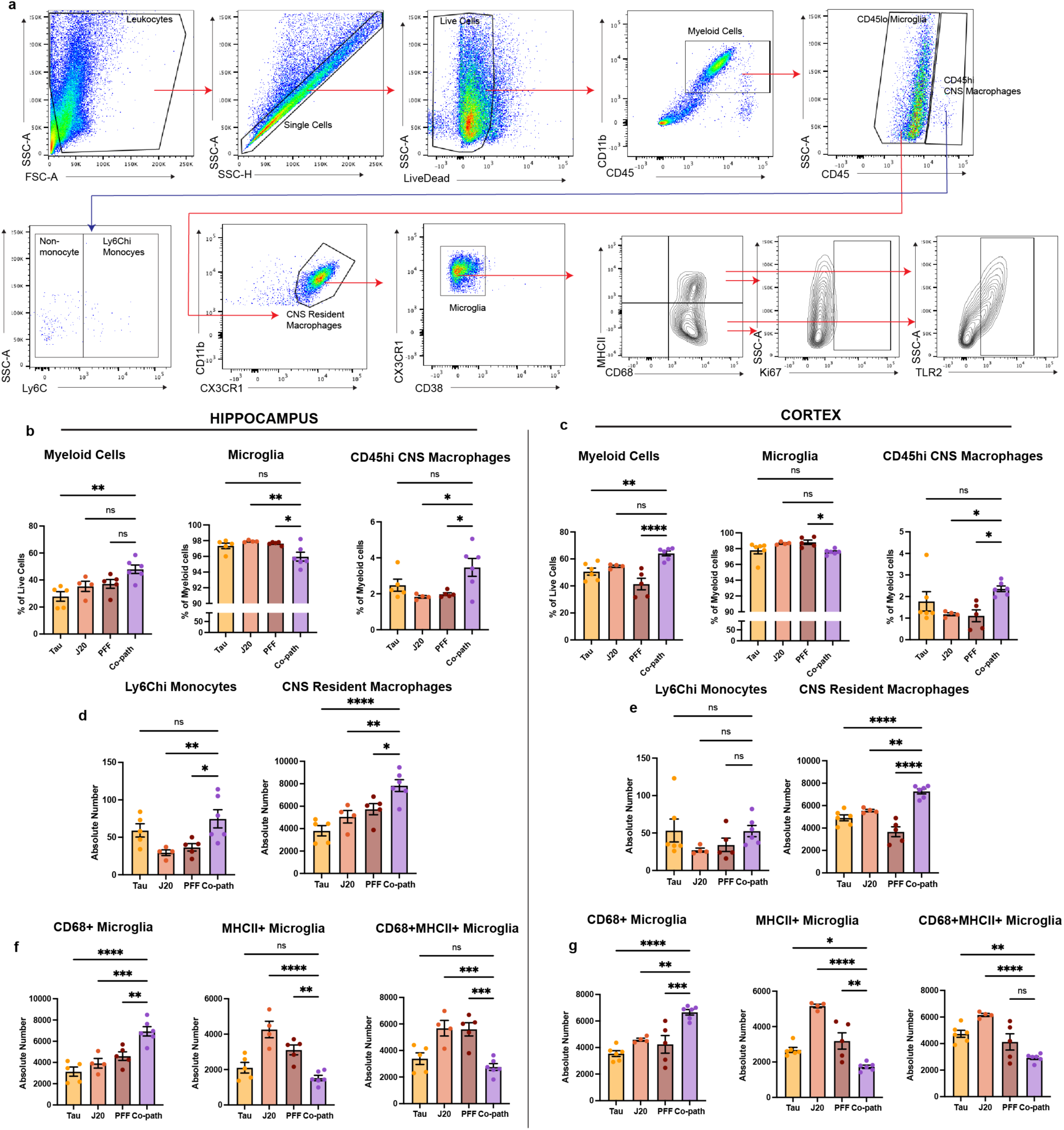
Myeloid cell flow cytometry gating, validation and quantification in co-pathology and single pathology animals. (a) Gating strategy for flow cytometry analysis of microglia. (b) Quantification of proportion of myeloid cells, microglia and infiltrating monocytes/CNS resident macrophages in the co-pathology model hippocampus and (c) cortex compared to single pathology brains. (d,e) absolute number of CD68+ microglia, MHCII+ microglia and MHCII+CD68+ microglia in the hippocampus and cortex of single pathology and co-pathology brains. Analyzed using One-way ANOVA. Mean values +/- SEM are plotted. *p<0.05, **p<0.01, ***p<0.005. n=5-6/group, with two brain regions pooled per sampled, both males and females included in analysis.

**Supplemental Figure 6.**
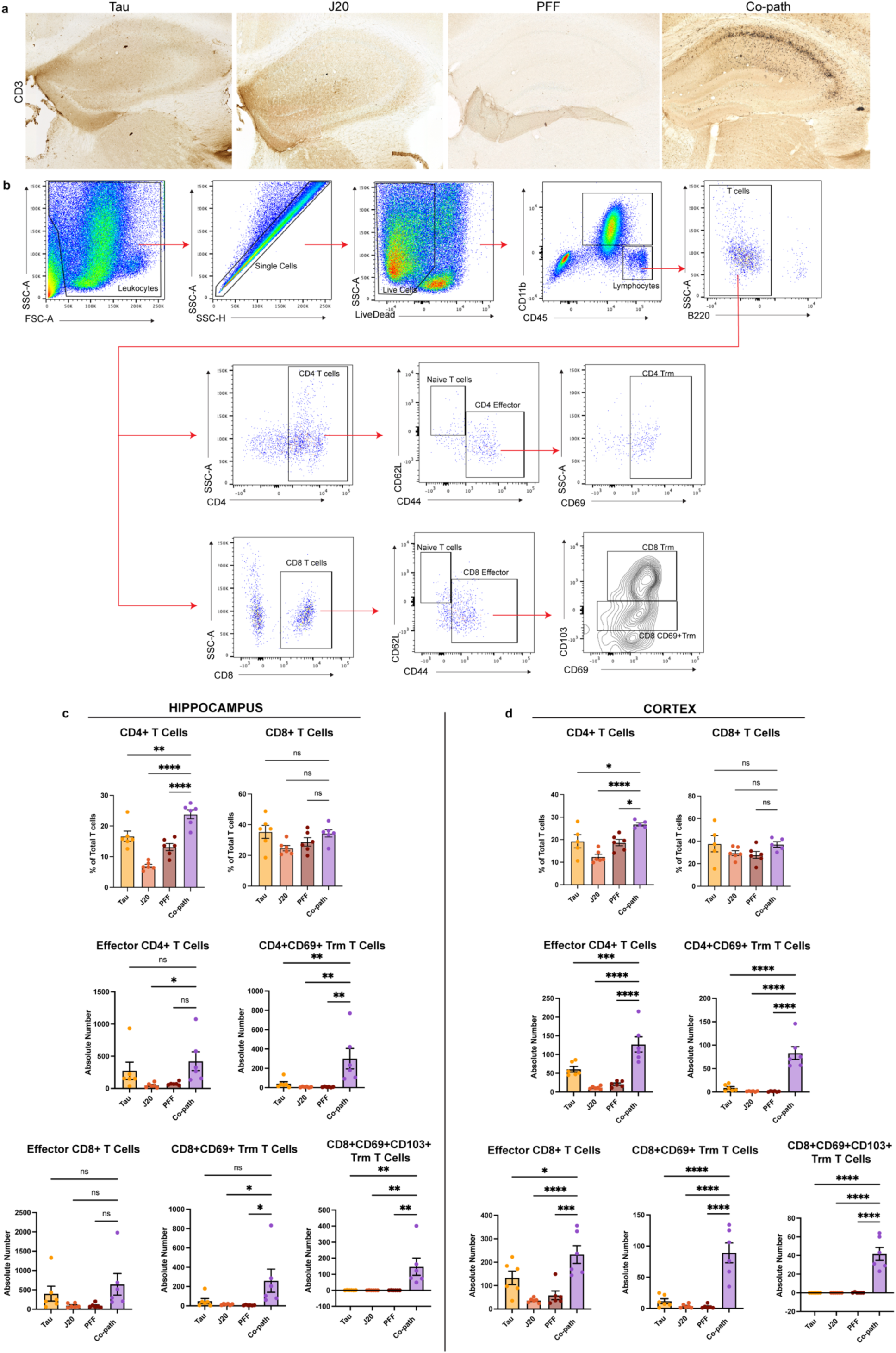
T cell flow cytometry gating, validation and quantification in co-pathology and single pathology animals. (a) Representative images showing CD3+ staining (nickel) from Tau, J20, PFF and co-pathology brains. (b) Gating strategy for flow cytometry analysis of T cells. (c) Quantification from flow cytometry of proportion of CD4 and CD8 T cells, and absolute number of effector CD4 and CD8 T cells, and Trm T cells in the hippocampus and (d) cortex. Analyzed using One-way ANOVA. Mean values +/- SEM are plotted. *p<0.05, **p<0.01, ***p<0.005. n=5-6/group, with two brain regions pooled per sampled, both males and females included in analysis.

